# The mycomembrane differentially and heterogeneously restricts antibiotic permeation

**DOI:** 10.1101/2024.12.31.630956

**Authors:** Irene Lepori, Kiserian Jackson, Zichen Liu, Mahendra D. Chordia, Mitchell Wong, Sylvia L. Rivera, Marta Roncetti, Laura Poliseno, Marcos M. Pires, M. Sloan Siegrist

## Abstract

The recalcitrance of *Mycobacterium tuberculosis* to antibiotic treatment has been broadly attributed to the impermeability of the organism’s outer mycomembrane. However, the studies that support this inference have been indirect and/or reliant on bulk population measurements. We previously developed the <u>P</u>eptidoglycan <u>A</u>ccessibility <u>C</u>lick-<u>M</u>ediated <u>A</u>ssessme<u>N</u>t (PAC-MAN) method to covalently trap azide-modified small molecules in the peptidoglycan cell wall of live mycobacteria, after they have traversed the mycomembrane. Using PAC-MAN we now show that the mycomembrane differentially restricts access of fluorophores and antibiotic derivatives. Mycomembranes of both *M. tuberculosis* and the model organism *M. smegmatis* discriminate between divergent classes of antibiotics as well as between antibiotics within a single family, the fluoroquinolones. By analyzing sub-populations of *M. tuberculosis* and *M. smegmatis*, we also found that some fluorophores and vancomycin are heterogeneously restricted by the mycomembrane. Our data indicate that the mycomembrane is a molecule- and cell-specific barrier to antibiotic permeation.

**Abstract Graphic:** 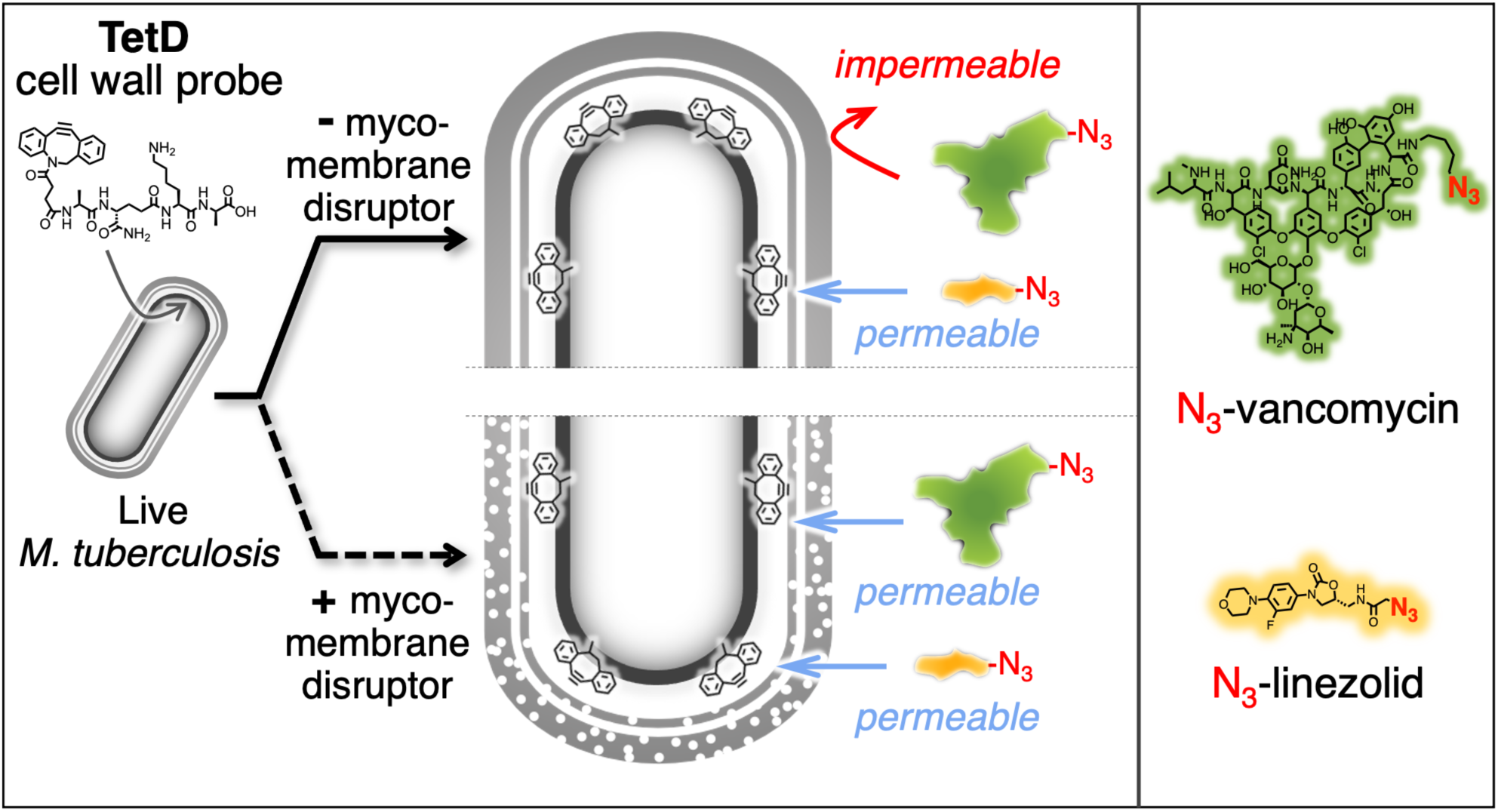

## Introduction

*Mycobacterium tuberculosis* infections cause approximately 1.5 million deaths annually^1^. The standard course of therapy for drug-susceptible tuberculosis is at least four months of four front-line antibiotics. Treatment for drug-resistant tuberculosis can take up to 2 years and relies on second-line antibiotics that are less effective, more toxic, and more expensive. Shortening the duration of treatment is a long-standing goal in the field.

In addition to host and host-induced determinants of antibiotic efficacy, *M. tuberculosis* is intrinsically less susceptible to drugs than monoderm bacteria that lack outer membranes^2–5^. High drug doses are often required for tuberculosis treatment, which in turn increase the risk of side effects and reduce compliance with the regimen^6^. The intrinsic resistance of *M. tuberculosis* is also a major hurdle for translating the results of *in vitro* target-based screening to the *in vivo* environment.

Encasing the cytoplasm, the mycobacterial cell envelope consists of an inner plasma membrane and peptidoglycan cell wall that is, in turn, covalently attached to a layer of arabinogalactan and an outer mycomembrane. Arabinogalactan-linked mycolates and noncovalent mycolates and other lipids respectively form the inner and outer leaflets of the mycomembrane, a distinctive feature of the *Corynebacterineae* suborder^2–5^. As mycomembrane-disrupting mutations and small molecules can sensitize mycobacteria to antibiotics and increase uptake of certain fluorescent dyes^7–13^ it is often presumed that the mycobacterial envelope, particularly the mycomembrane, is a key determinant of impermeability and therefore intrinsic antibiotic resistance.

There are also hints in the literature that the mycobacterial envelope may pose a differential and heterogenous barrier to permeation. Genetic disruption of the envelope sensitizes *M. tuberculosis* to vancomycin, rifampicin, and bedaquiline but not to other antibiotics tested^13^. As well, the model organism *M. smegmatis* exhibits single-cell heterogeneity in susceptibility to some antibiotics that correlates with heterogeneity in fluorescent dye uptake^14–16^. The nature of envelope restriction has implications for drug development. For example, a differential and/or heterogeneous barrier could indicate that adjuvants that disrupt the envelope are not a one-size-fits-all strategy to potentiate antibiotics for tuberculosis.

Here we applied our recently-developed assay, <u>P</u>eptidoglycan <u>A</u>ccessibility <u>C</u>lick-<u>M</u>ediated <u>A</u>ssessme<u>N</u>t (PAC-MAN^17^), to test long-standing inferences related to mycomembrane permeation. PAC-MAN uses metabolic labeling and bioorthogonal click chemistry to covalently trap azide-modified small molecules in the cell walls of live bacteria. By querying cell wall accessibility, rather than whole cell accumulation, we aimed to interrogate molecule permeation in close proximity to the mycomembrane. We also aimed to gauge permeation at both the population and single-cell levels by leveraging PAC-MAN’s flow cytometry readout. Using *M. tuberculosis* and the model organism *M. smegmatis*, we found that the mycomembrane differentially restricts periplasm access of a diverse set of fluorophores and antibiotic derivatives. The mycomembrane differentiates between divergent classes of antibiotics, as anticipated from activity studies, as well as between molecules within a single family, the fluoroquinolones. We also found that some fluorophores and vancomycin are heterogeneously restricted by the mycomembrane. The complexity and variability of the mycomembrane permeability barrier suggest the utility of molecule-specific strategies to breach it.

## Results

### Diderm outer membranes differentially restrict fluorophores

We and others developed D-amino acid-based probes that incorporate at sites of active cell wall metabolism in bacteria^18–25^. A subset of these probes is directly conjugated to fluorophores. In the model Gram-negative bacterium *Escherichia coli*, incorporation of a D-amino acid conjugated to 7-hydroxycoumarin is less sensitive to outer membrane perturbation compared to D-amino acids conjugated to other dyes^26^. These data suggest that the *E. coli* outer membrane differentially restricts fluorophore access to the periplasm.

We sought to test whether the mycomembrane is similarly a differential barrier. However, we previously found in *M. smegmatis* that the structure of the fluorophore conjugated to the D-amino acid can bias the probe’s enzyme-mediated incorporation route^27^. Therefore, we took advantage of a two-step labeling process whereby we label the cell wall using a strained cyclooctyne-conjugated D-amino acid probe^21^ and detect via strain-promoted alkyne-azide cycloaddition (SPAAC^28–30^; **Figure 1A**) with diverse azide-modified fluorophores (**Supplemental Figure 1** and **Supplemental Table 1**). By separating the probe incorporation step from the fluorophore label, our goal was to isolate fluorophore structure as the primary variable for mycomembrane permeation.

**Figure 1.**
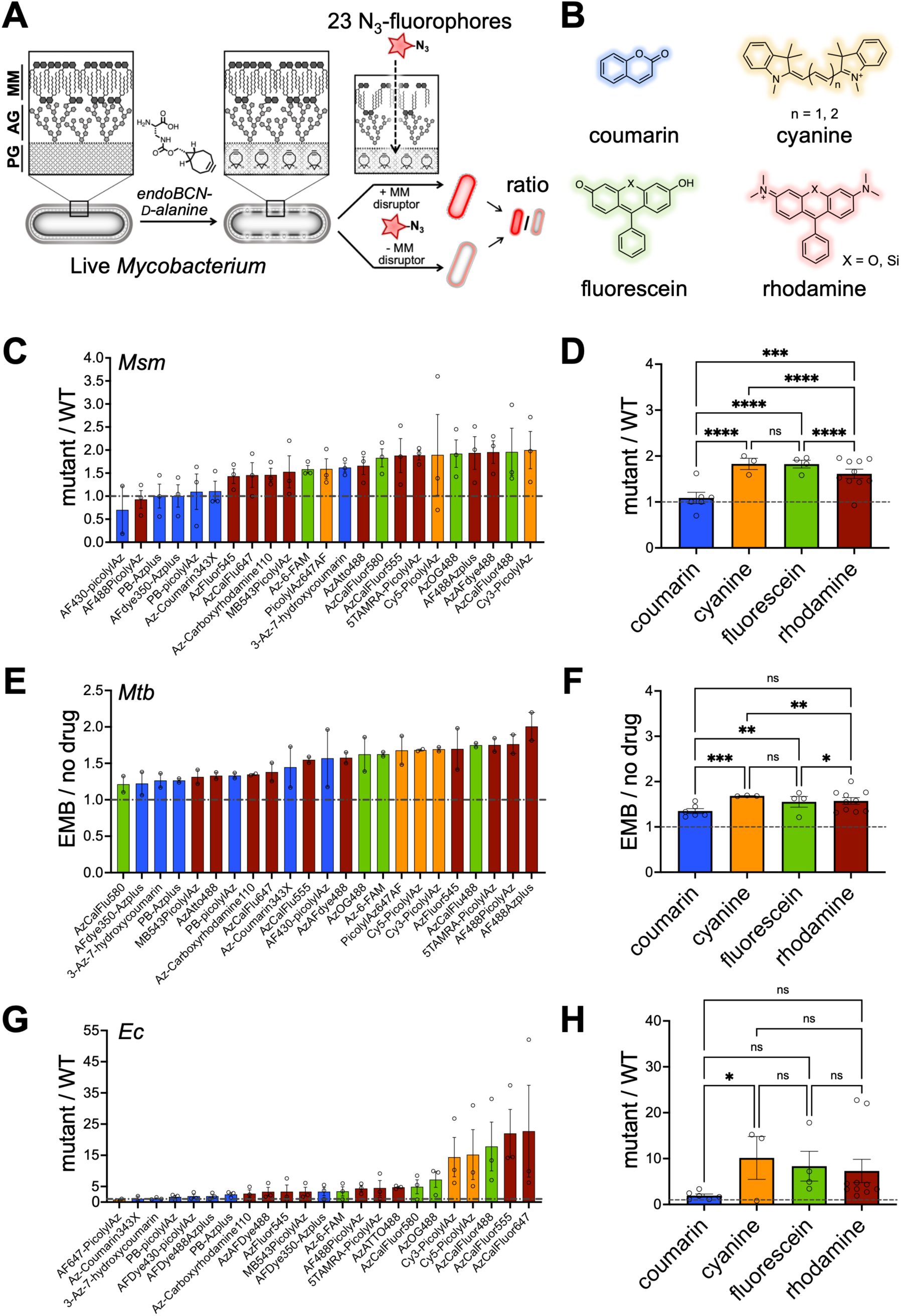
Diderm outer membranes differentially restrict fluorophores. (A) Schematic representation of the experimental workflow used in (C-H). Bacteria with or without a mycomembrane-disrupting mutation are incubated with BCN-_D_-alanine (BCNDala) probe, to tag the peptidoglycan, then subjected to SPAAC reaction with different azide fluorophores (**Supplemental Figure 1**). For chemical perturbation of the mycomembrane, bacteria were treated with drug after BCNDala and before SPAAC. Fluorescence plotted as a ratio between values from bacteria with disrupted outer (myco)membranes vs. bacteria with undisrupted outer (myco)membranes. PG: peptidoglycan; AG: arabinogalactan; MM: mycomembrane. (B) Chemical scaffolds of the four investigated dye families: coumarin, cyanine, fluorescein, and rhodamine. *M. smegmatis* (C), *M. tuberculosis* (E), and *E. coli* (G) cell wall were treated as shown in (A) and the fluorescence ratio of outer (myco)membrane-disrupted (*M. smegmatis* Δ*fbpA*, *M. tuberculosis* + ethambutol (EMB), *E. coli lptD*4213) vs. undisrupted (*M. smegmatis* wild-type, *M. tuberculosis* no drug, *E. coli* wild-type) were determined. Results from (C), (G), and (E) were graphed by fluorophore scaffold in (D), (F), and (H). Coumarins change less with outer (myco)membrane perturbation compared to the other fluorophore scaffolds. Data are represented as mean +/− SEM of 2-3 independent biological replicates. *Msm: M. smegmatis; Mtb: M. tuberculosis, Ec: E. coli.* *, **, ***, and ****, p < 0.05, 0.005, 0.0005, and 0.00005 respectively by two-way ANOVA on log_2_-transformed data.

After labeling the cell walls of live *M. smegmatis* and human pathogen *M. tuberculosis* with a D-amino acid conjugated to bicyclo[6.1.0]non-4-yne (BCNDala^21^), we exposed the bacteria to a panel of commercial azide fluorophores. The labels varied in chemical scaffold, expected azide reactivity, presence or absence of a linker, charge, fluorogenicity, among other features (**Supplemental Figure 1**). To test the role of mycomembrane integrity, we compared wild-type to Δ*fbpA M. smegmatis* and untreated to ethambutol-treated *M. tuberculosis*. Loss of FbpA (Antigen 85A) directly disrupts the mycomembrane, by inhibiting the transfer of mycolic acids to the inner and/or outer leaflets, while ethambutol indirectly disrupts the mycomembrane by inhibiting biosynthesis of the underlying arabinogalactan^5, 31–35^. As anticipated, fluorescence quantitated by flow cytometry generally increased in Δ*fbpA M. smegmatis* relative to wild-type and in ethambutol-treated *M. tuberculosis* relative to untreated (**Figure 1C, 1E**). As a control, we performed a similar analysis in *E. coli,* comparing wild-type to an *lptD*4213 strain^36, 37^. *E. coli* with the *lptD*4213 allele have a compromised outer membrane barrier^38^. Similar to mycobacterial species, fluorescence generally increased upon *E. coli* outer membrane perturbation (**Figure 1G**).

To test whether the mycomembrane is a differential barrier to fluorophores, we compared the effect of mycomembrane perturbation on labeling by four dye families: coumarin, cyanine, fluorescein, and rhodamine (**Figure 1B**, **Supplemental Figure 1**). We were able to discern scaffold-specific differences (**Figure 1D, 1F, 1H**); other physicochemical properties and features of the molecules did not obviously impact mycomembrane permeation (**Supplemental Figure 2**). Specifically, we found that coumarin derivatives are relatively insensitive to *fbpA* deletion or ethambutol treatment compared to cyanine, fluorescein, and rhodamine derivatives. Likewise, coumarin derivatives are less sensitive to *E. coli lptD* mutation compared to cyanines, and trended similarly compared to fluorescein and rhodamine. Taken together, we conclude that diderm outer (myco)membranes restrict fluorophore access to the periplasm in scaffold-specific manner.

### Fluorophore restriction by the mycomembrane is heterogeneous

Mycobacterial species grow and divide heterogeneously, a property that has been linked to differential labeling by a fluorescent probe as well as differential susceptibility to rifampicin and some envelope-acting drugs^14–16^. Upon closer inspection of the flow cytometry data, we found that *M. smegmatis* and *M. tuberculosis* labeled relatively homogeneously when exposed to coumarin derivatives but often comprised either multiple populations or much broader distributions when exposed to cyanine, fluorescein, or rhodamine derivatives (**Figure 2A**). This phenotype was more pronounced in *M. tuberculosi*s. The proportion of cells that were labeled was inversely proportional to optical density of the culture and increased substantially upon ethambutol treatment (**Figure 2B-D**). These data suggest that there is heterogeneity in mycomembrane restriction of fluorophores.

**Figure 2.**
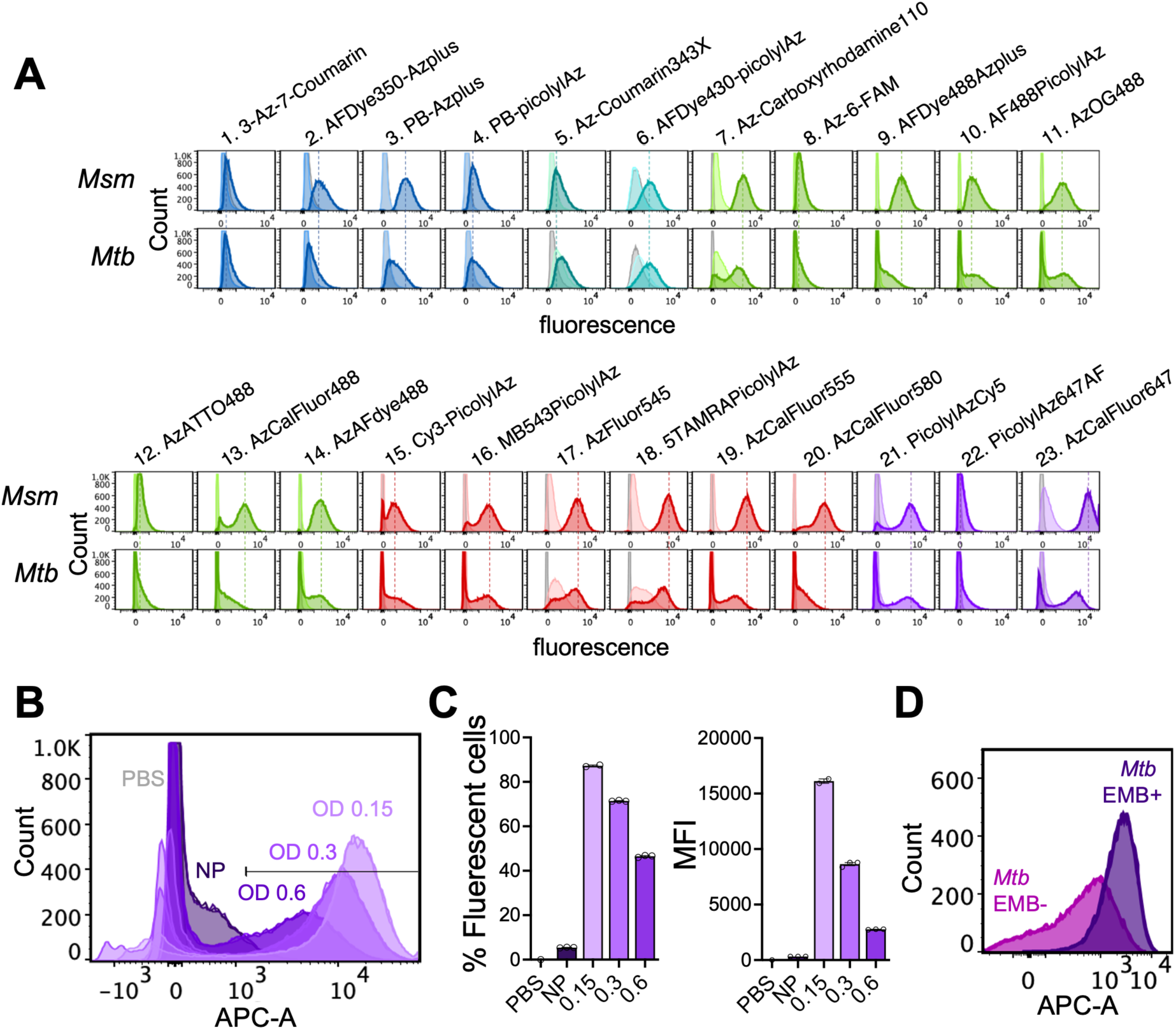
Fluorophore restriction by the mycomembrane is heterogeneous. (A) Flow cytometry histograms for the labeling of *M. smegmatis* and *M. tuberculosis* with azide fluorophores (**Supplemental Figure 1**), using **TetD** to tag the peptidoglycan. *M. tuberculosis* comprises two populations for all fluorophores except for the coumarin derivatives. *M. smegmatis* also comprises two populations for some non-coumarin fluorophores. (B). Flow cytometry histograms for the labeling of *M. tuberculosis* with AzCalFluor467. Bacteria were taken from the indicated growing OD_600_ then normalized to the same working OD_600_ prior to fluorophore exposure. (C) Both the percentage of fluorescent cells and the mean fluorescence intensities (MFI) are inversely proportional to the original, growing OD_600_. (D) Ethambutol (EMB) pre-treatment results in more uniform labeling. Histograms and the graphs are representative of 3 independent biological replicates. Data in (C) and (D) are mean+/− SEM of three technical replicates. *Msm: M. smegmatis; Mtb: M. tuberculosis*.

### The mycomembrane differentially restricts antibiotic activity and permeation

Our fluorophore data suggested that the mycomembrane is a differential barrier to permeation of fluorophore scaffolds. We next wished to test whether this observation extended to antibiotics. Outer membrane perturbation has been found to sensitize diderm bacteria to vancomycin and rifampicin (*M. tuberculosis* and Gram-negative species) and novobiocin (*M. smegmatis* and Gram-negatives) but not to linezolid (*M. tuberculosis*)^11, 13, 39–41^. We first verified in *M. smegmatis* that mycomembrane perturbation via *fbpA* deletion sensitizes the bacteria to killing by vancomycin, rifampicin, and novobiocin but not to linezolid (**Figure 3A**). We also verified that mycomembrane perturbation via ethambutol sensitizes *M. tuberculosis* to novobiocin (**Figure 3B**).

**Figure 3.**
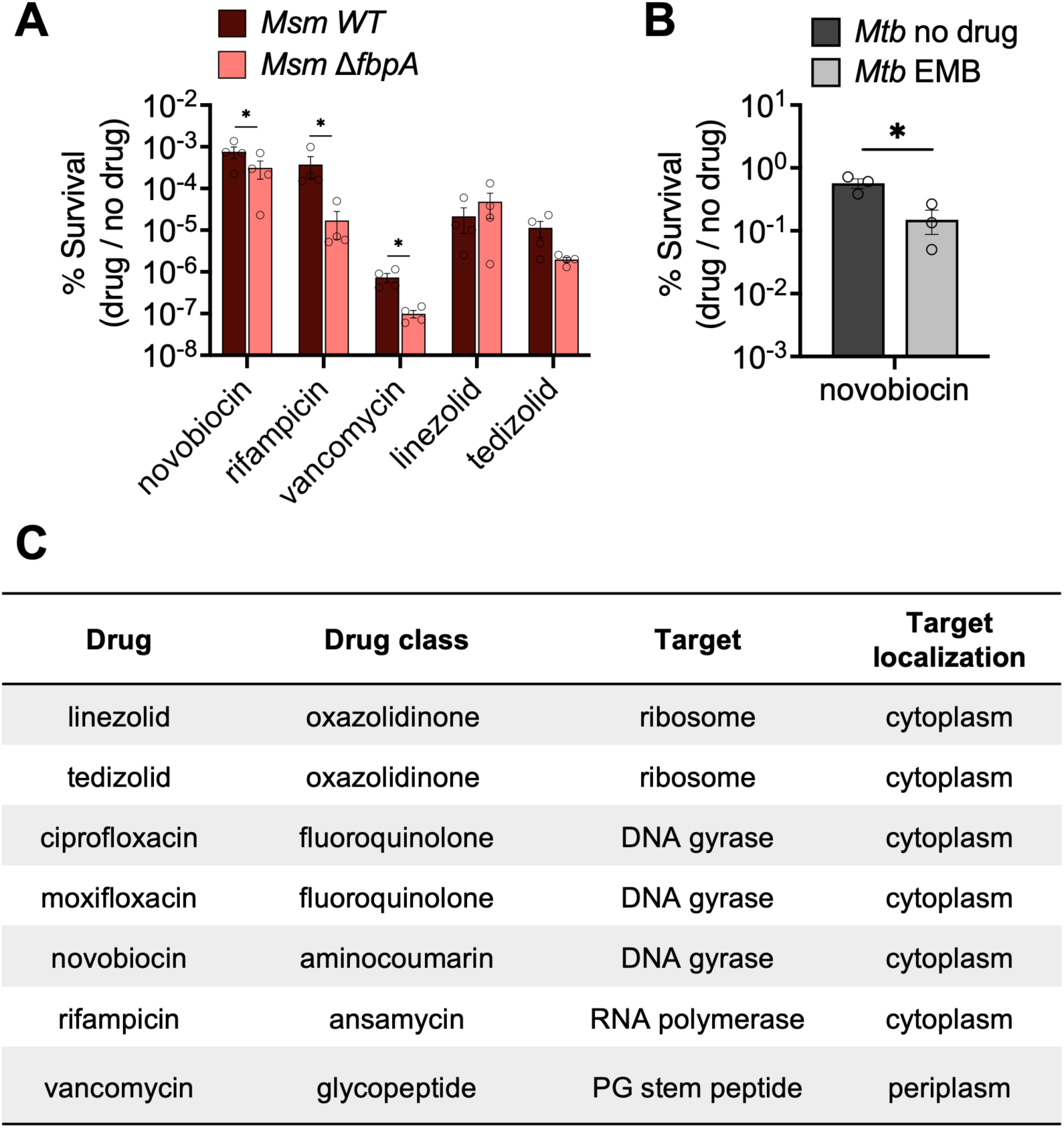
The *M. smegmatis* mycomembrane differentially restricts antibiotic activity. (A) Loss of Antigen 85A (Δ*fbpA*) sensitizes *M. smegmatis* to novobiocin, rifampicin, and vancomycin, but not to linezolid and tedizolid. (B) Novobiocin is more effective against *M. tuberculosis* pre-treated with ethambutol (EMB). Bacteria were exposed to 50 *μ*M drug, to match subsequent azide antibiotics experiments, and plated for colony forming units (CFU) after 24 hours. Survival plotted as a ratio of drug-treated to untreated. (C) Antibiotics investigated in this work and related features. *Msm: M. smegmatis; Mtb: M. tuberculosis*. PG: peptidoglycan. Data are represented as mean +/− SEM of 3-4 independent biological replicates. *, p < 0.05 by paired t-test on log_10_ transformed data.

We hypothesized that the differential activity of these antibiotics +/− *fbpA* or +/− ethambutol reflects differential permeation across the mycomembrane. We used our previously-developed, three-step PAC-MAN assay (**Figure 4A**) for measuring accumulation of non-fluorescent molecules^17^ wherein we label the cell walls of live mycobacteria with a strained cyclooctyne-bearing probe, as above (*Step 1*; here we use a tetrapeptide peptidoglycan stem peptide mimic that bears a dibenzocyclooctyne (DBCO) reactive group (**TetD**^17^)); pulse with an azide-modified test molecule of interest (*Step 2*; test azide); and chase with an azide fluorophore (*Step 3*). Because *Step 1* and *Step 3* are constant, and we normalize to controls for which individual steps are omitted, perturbations that impact **TetD** incorporation and/or azide fluorophore accessibility are controlled within a given condition. Indeed, we found that small (2-5-fold) changes in **TetD** or azide fluorophore concentration do not significantly affect PAC-MAN results, nor does choice of azide fluorophore, the latter despite the heterogeneity in labeling noted above (**Supplemental Figure 3**).

**Figure 4.**
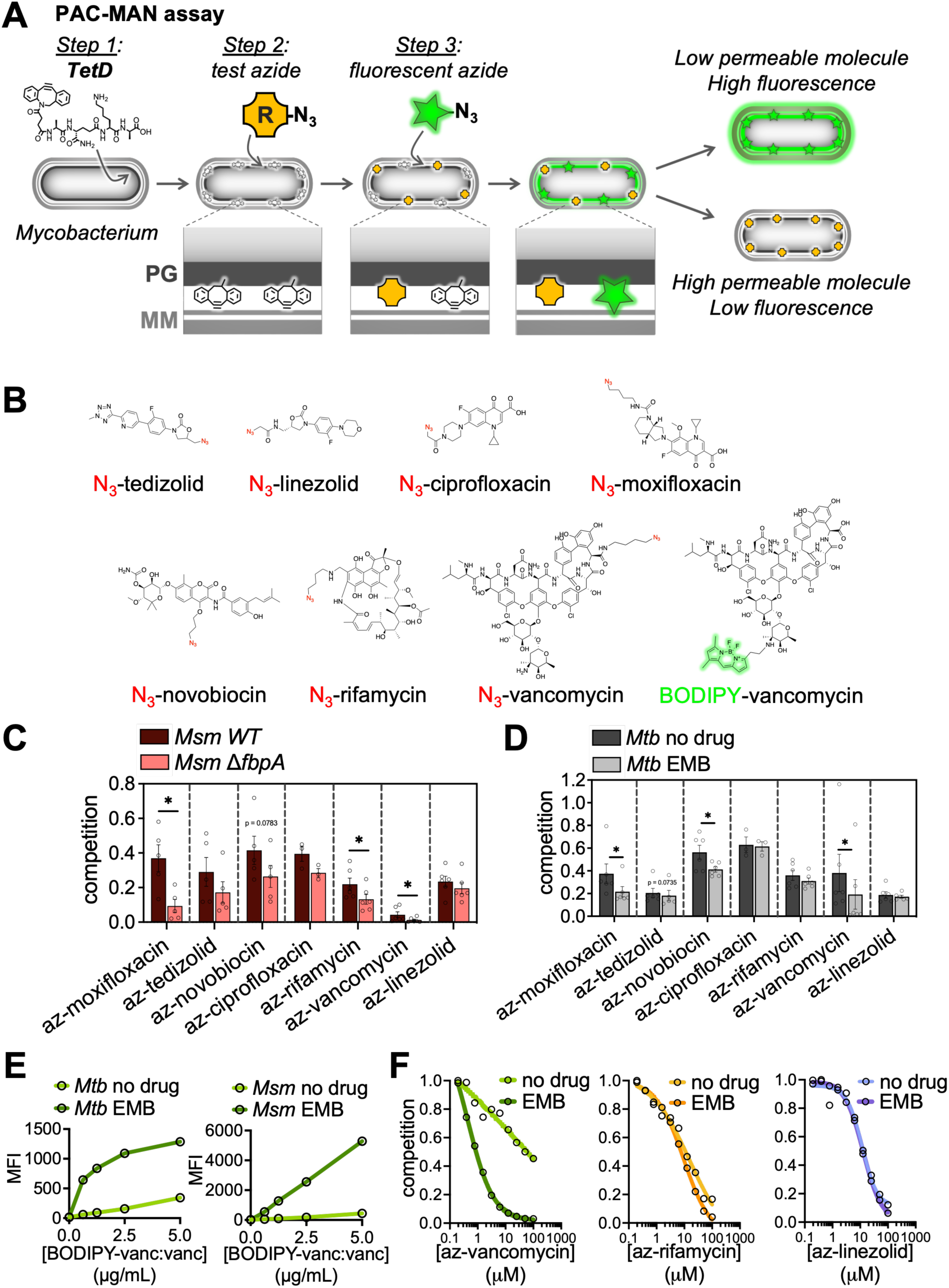
The mycomembrane differentially restricts antibiotic permeation. (A) Schematic of PAC-MAN assay. In *Step 1*, bacteria are incubated with the **TetD** probe to tag the peptidoglycan. Then, bacteria are incubated with an azide-modified test molecule (*Step 2*) followed by an azide fluorophore (*Step 3*). Test azides that do not permeate the mycomembrane do not access cell wall-embedded DBCO. DBCO that do not react in *Step 2* are free to react with azide fluorophores in *Step 3*, resulting in high fluorescence. Conversely, test azides that permeate well can access DBCO in *Step 2*, leaving fewer unreacted DBCO to react with azide fluorophores in *Step 3* and resulting in low fluorescence. (B) Chemical structures of azide (az) antibiotics used in this work and BODIPY-vancomycin. (C-D) PAC-MAN assay for (C) *M. smegmatis* wild-type and Δ*fbpA* or (D) *M. tuberculosis* +/− ethambutol (EMB) using the azide antibiotics in (B). Permeation of vancomycin and moxifloxacin significantly increases upon mycomembrane perturbation for both species; rifamycin increases in *M. smegmatis*; novobiocin increases in *M. tuberculosis* and trends similarly in *M. smegmatis*. Competition levels of azide antibiotics in (C) and (D) are comparable between strains, treatments, and species but not to each other due to differences in azide antibiotic reactivity. See Figure 6 and accompanying text for details. (E) BODIPY-vancomycin labeling of *M. smegmatis* and *M. tuberculosis* increases with concentration and upon EMB pre-treatment. (F) Competition in *M. tuberculosis* +/− EMB for different concentrations of azide-vancomycin, -rifamycin, and -linezolid. The three antibiotics respectively show high, slight, and no sensitivity to mycomembrane disruption. *Msm: M. smegmatis; Mtb: M. tuberculosis*. *, p<0.05 by paired t-test performed on log_10_-transformed data.

To test for differential restriction of antibiotics across the mycomembrane, we compared accumulation of azide-modified antibiotics^42^ (**Figure 4B**) in wild-type, Δ*fbpA*, and complemented Δ*fbpA M. smegmatis* as well as untreated and ethambutol-treated *M. tuberculosis*. Mycomembrane perturbation enhanced azide-vancomycin accumulation and trended the same for azide-novobiocin accumulation (**Figure 4C-D** and **Supplemental Figure 4**). We recapitulated the vancomycin difference using a fluorescent conjugate of the drug (**Figure 4E**). While mycomembrane perturbation enhanced azide-rifamycin accumulation in *M. smegmatis*, consistent with activity data (^13^ and **Figure 3A**), the effect in *M. tuberculosis* was not statistically significant. We reasoned that we might be able to capture more subtle differences by querying the azide antibiotics across a range of concentrations. Using this protocol, we verified that ethambutol treatment enhanced the accumulation of azide-vancomycin to a much greater degree than azide-rifamycin in *M. tuberculosis* and had no effect on azide-linezolid (**Figure 4F**).

The effect of mycomembrane perturbation on azide-rifamycin accumulation appears more modest than that observed previously for a fluorescein conjugate of rifampicin^32^. However, given our observation that fluorescein-based dyes are themselves sensitive to *fbpA* deletion and ethambutol in *M. smegmatis* and *M. tuberculosis*, respectively, we speculated that our results were more consistent with more marginal increases observed for [14C] rifampicin accumulation upon ethambutol treatment of *M. smegmatis* or *M. tuberculosis*^43^. In principle, metabolic transformation of azide antibiotics may decrease the number of azides available for reaction with DBCO, limiting our dynamic range and making it more difficult to assess changes in the presence or absence of mycomembrane perturbation. However, the time frame (two hours exposure to azide antibiotics) and cellular compartment (periplasm rather than cytoplasm, where known xenobiotic metabolism enzymes reside) suggested an alternative or additional explanation. Other groups have noted connections between rifampicin treatment and/or resistance and cell envelope synthesis and/or stress, both in mycobacterial species and other organisms^8, 44–50^. Thus, we considered whether synergism between rifampicin and mycomembrane perturbation could be due at least in part to an effect of rifampicin on the envelope. Indeed, we find that while cell wall metabolism is robust in rifampicin-treated *M. smegmatis*, it is delocalized from the normal sites of polar cell elongation (**Supplemental Figure 5**). Although not definitive, these data support the notion of a bidirectional synergy between perturbations to transcription and the mycobacterial envelope.

### The mycomembrane is a more restrictive barrier for moxifloxacin than for ciprofloxacin

Vancomycin, rifampicin, novobiocin, and linezolid comprise distinct classes of antibiotics (**Figure 3C**). We wondered whether the mycomembane might also differentially restrict antibiotics that belong to the same class, *i.e.*, share a target and have similar structures. Moxifloxacin has better activity against mycobacterial species compared to the structurally-related fluoroquinolone ciprofloxacin^51–55^ despite inhibiting DNA gyrase to similar degrees *in vitro*^56^. Because the whole cell anti-mycobacterial activities of fluoroquinolones correlate with their hydrophobicities^51^, it was previously hypothesized that moxifloxacin permeates the mycomembrane more readily than more hydrophilic fluoroquinolones like ciprofloxacin^53^. Surprisingly, we found that mycomembrane perturbation enhanced azide-moxifloxacin accumulation more than azide-ciprofloxacin (**Figure 4C-D**). These data suggest that the mycomembrane restricts the more-hydrophobic moxifloxacin to a greater extent than ciprofloxacin.

We also compared the effect of ethambutol and/or *fbpA* mutation on two oxazolidinones. While mycomembrane perturbation modestly impacted the activity of tedizolid more than linezolid (**Figure 3A**), and the accumulation of azide-tedizolid more than azide-linezolid, the effects were not statistically significant (**Figure 4C-D**).

### Vancomycin restriction by the mycomembrane is heterogeneous

We previously observed that *M. smegmatis* labeled with a BODIPY conjugate of vancomycin comprised fluorescent and non-fluorescent populations^27^. We verified by flow cytometry that this observation held for both *M. smegmatis* and *M. tuberculosis* (**Figure 5A**), and further that ethambutol treatment resulted in labeling that was both brighter and more uniform in its distribution. To test whether homogenization of BODIPY-vancomycin labeling by mycomembrane disruption reflected changes in the accumulation of drug, the dye, or the combination, we asked whether azide-vancomycin behaved similarly. We opted to focus on *M. smegmatis* for this experiment, as its azide fluorophore labeling at baseline (in the absence of azide antibiotic in *Step 2*; **Figure 4A**), is more homogenous than *M. tuberculosis* (**Figure 2A**; azide fluorophore #23) and therefore more straightforward to interpret. We reasoned that heterogenous incorporation of an azide antibiotic in *Step 2* of our assay should result in unequal cell-to-cell distribution of unreacted DBCO moieties available for detection by azide fluorophore in *Step 3* (**Figure 4A**). We found that while azide-vancomycin decreased overall fluorescence of *M. smegmatis* compared to controls lacking the azide-vancomycin competitor, it also increased the variance of the population, suggesting that it incorporates heterogeneously (**Figure 5B**). We did not reproducibly observe this phenotype for any of the other drug derivatives we tested (**Supplemental Figure 6**).

**Figure 5.**
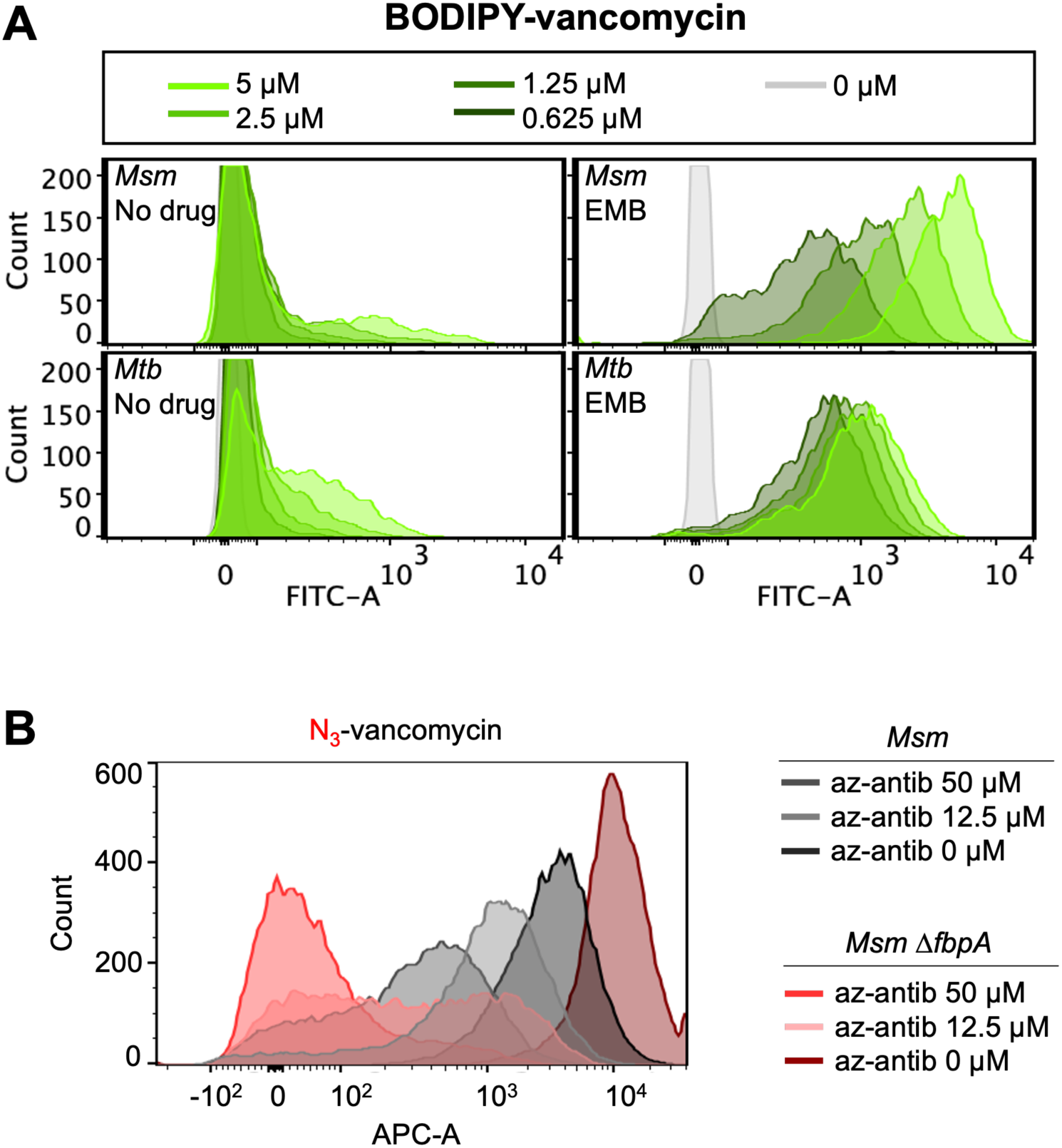
Vancomycin restriction by the mycomembrane is heterogeneous. A) *M. smegmatis* (top) and *M. tuberculosis* (bottom) were pre-treated or not with ethambutol (EMB) and exposed to different concentrations of BODIPY-vancomycin. Both organisms exhibit heterogeneity, with an unlabeled population (left panels) that decreases with upon ethambutol pre-treatment (right panels). (B) Flow cytometry histograms from PAC-MAN for azide-vancomycin*. M. smegmatis* exhibits variable competition, most evident at 12.5 *μ*M for Δ*fbpA* and 50 *μ*M for wild-type.

We hypothesized that the heterogeneity in azide-vancomycin incorporation reflected heterogeneity in mycomembrane permeation. However, azide-vancomycin incorporation remained heterogeneous in the absence of *fbpA* (**Figure 5B**), a surprising result given that mycomembrane disruption by ethambutol had resulted in more uniform labeling by BODIPY-vancomycin (**Figure 5A**). We speculate that there are different sources of heterogeneity within the mycobacterial envelope, and further that envelope components have molecule-specific effects on permeation. Nevertheless, our data suggest that the mycomembrane heterogeneously restricts the accumulation of vancomycin.

### Differential restriction of cytoplasm-acting antibiotics by the mycomembrane correlates with periplasm accumulation

We wished to probe the relationship between mycomembrane permeation and periplasm accumulation. However, our analyses above, while appropriate for cross-perturbation or cross-species comparisons, did not take into account the intrinsic reactivity of the test azide. Each test azide varies in both reactivity and permeation; we wished to isolate the latter variable for cross-azide comparisons

We previously addressed this question by reacting **TetD**-labeled mycobacteria in parallel with DBCO-bearing polystyrene beads, the latter of which lack a membrane barrier to restrict access of the test azide to its cyclooctyne partner. We then analyzed only those test azides above a specific threshold of reactivity^17, 42^. To broaden the scope of analyzable test azides, we developed a new protocol for normalization inspired by the chloroalkane penetration assay (CAPA) developed by the Kritzer group^57^. In the CAPA approach, cells expressing the HaloTag enzyme are pulsed with different concentrations of a test chloroalkane (covalent ligand for HaloTag), washed, then challenged with a fluorescent chloroalkane to detect HaloTags that did not react with the test chloroalkane in the step prior. The extent of test chloroalkane accumulation is then calculated as the concentration that results in 50% reduction in fluorescence. Similarly, we exposed DBCO-bearing polystyrene beads and **TetD**-labeled *M. tuberculosis* to different concentrations of azide antibiotics and calculated the <u>c</u>oncentration of azide antibiotic that <u>c</u>ompetes fluorescence by <u>50</u>%, or “CC_50_” (**Figure 6A**). We reasoned that the difference in bead CC_50_ and bacterial CC_50_ for a given azide antibiotic should report relative periplasm accumulation.

**Figure 6.**
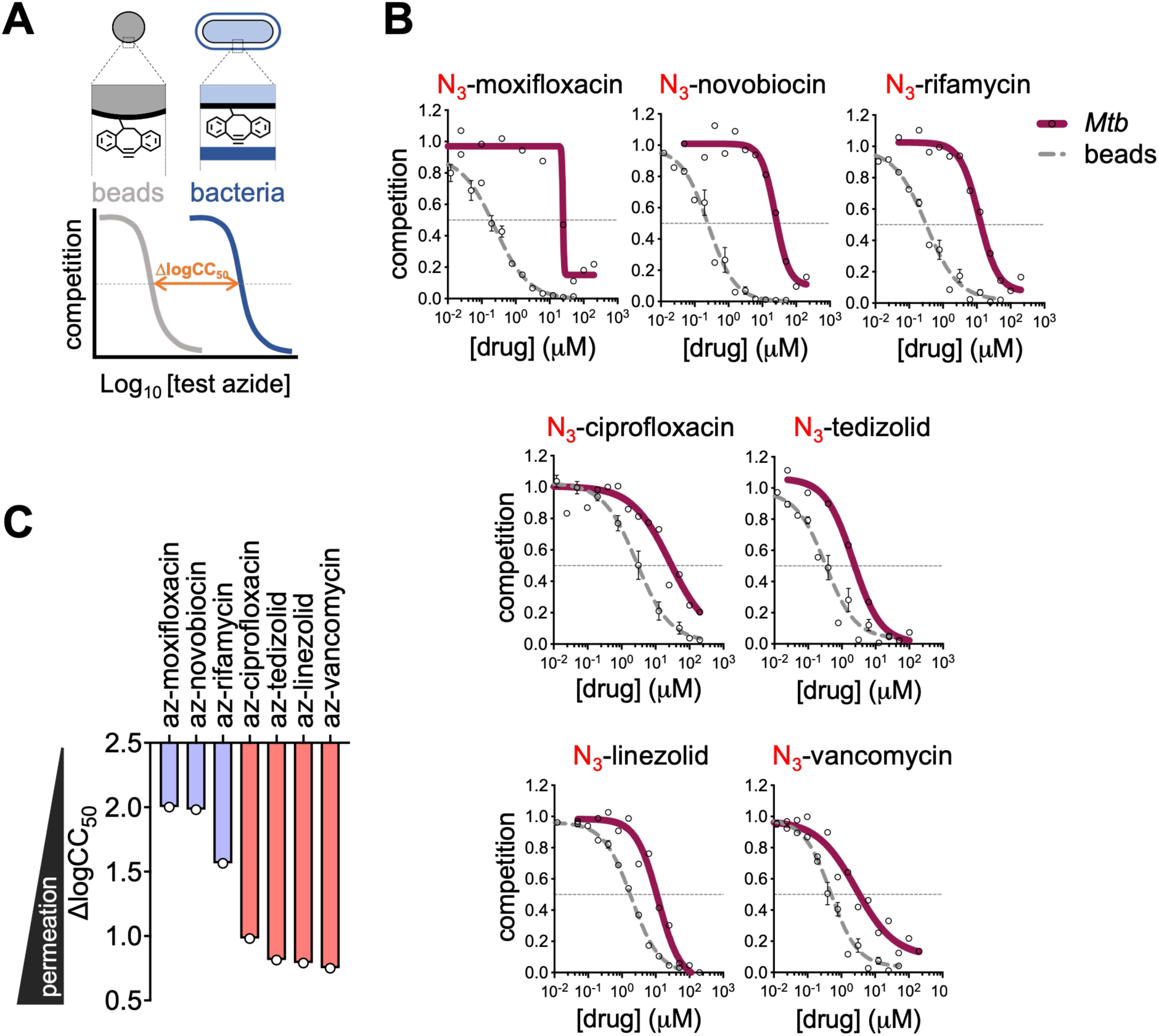
Differential accumulation of antibiotics in *M. tuberculosis* periplasm. To control for intrinsic differences in azide antibiotic reactivity, we performed PAC-MAN using different concentrations of azide antibiotics and compared the results from DBCO-modified polystyrene beads to **TetD**-labeled *M. tuberculosis* (A). Specifically, we calculated the <u>c</u>oncentration of azide drug that <u>c</u>ompetes fluorescence by <u>50</u>%, or “CC_50_,” and subtracted the log_10_CC_50_ on beads from log_10_CC_50_ on bacteria. (B) Competition for individual azide drugs on both DBCO-polystyrene beads and -*M. tuberculosis*. (C) Δlog_10_CC_50_ calculated from (B). Red and purple respectively highlight high- and low-accumulating drugs. Δlog_10_CC_50_ were calculated from four independent biological replicates for DBCO-polystyrene beads and at least two independent biological replicates for **TetD**-labeled *M. tuberculosis*.

For drugs with cytoplasmic targets, periplasm accumulation of their azide derivative corresponded with sensitivity to mycomembrane perturbation: azide antibiotics that were not significantly impacted by *fbpA* mutation in *M. smegmatis* or ethambutol treatment in *M. tuberculosis* (linezolid, tedizolid, ciprofloxacin; **Figure 4C-D**) generally accumulated to high levels in the periplasm (**Figure 6B-C**) whereas those that were significantly impacted by mycomembrane perturbation (novobiocin, moxifloxacin, and rifamycin; **Figure 4C-D**) accumulated to lower levels (**Figure 6B-C**). Vancomycin, which binds to cell wall and cell wall precursors and therefore acts in the periplasm, behaved differently: while the azide-modified drug was sensitive to both *fbpA* mutation and ethambutol treatment, it nevertheless accumulated to high levels in the periplasm (**Figure 6B-C**). The observation that azide-vancomycin is restricted by the mycomembrane but nonetheless accumulates relatively well in the periplasm is consistent with the location of its target^58–60^. It is also possible that the pre-SPAAC, non-covalent interaction between azide-vancomycin and cell wall (precursors) shifts the SPAAC reaction equilibrium to favor azide-vancomycin reaction with cell wall-embedded DBCO. Nonetheless, for the cytoplasm-acting antibiotics tested here, differential mycomembrane restriction is associated with differential accumulation.

## Discussion

There is a rich literature reporting the connection between mycomembrane alterations and sensitivity to a subset of antibiotics or fluorescent dyes^7–13^. These observations suggest that the mycomembrane is a key barrier to antibiotic accumulation and, therefore, whole cell activity. However antibiotic activity is an indirect proxy for uptake; for example, collateral metabolic dysfunction from cell envelope perturbation^50, 61–63^ may also impact drug sensitivity. Moreover, the chemical structures of fluorescent dyes used to report permeability, *e.g*., ethidium bromide, do not resemble those of antibiotics.

To address the role of the mycomembrane in antibiotic permeation, we used PAC-MAN^17^ to compare the relative accumulation of azide antibiotics in the periplasm, the cellular compartment bounded by the inner plasma membrane and mycomembrane. While we cannot rule out a potential effect of the azide modification on molecule permeation, we note that azides are widely recognized as optimal bioorthogonal tags due to their minimal, 3-atom size and typically-low impact on the physicochemical properties of the parent compound^28, 64^. We reasoned that accumulation of an azide antibiotic in the cell wall reflects the ability of a molecule to permeate across the mycomembrane and arabinogalactan layers of the envelope. PAC-MAN enables comparisons across species and across genetic and chemical perturbations, and by developing a new method for analysis, we were able to expand the utility of the assay to comparisons across azide-modified molecules. We find that the mycomembrane differentially restricts the periplasm accumulation of antibiotics and that this generally correlates with restriction of activity.

Beyond testing long-standing hypotheses about antibiotic permeation from the literature, our work adds nuance to the field in several ways. For example, synergism between envelope perturbations and rifampicin has frequently been attributed to permeabilization by the former enhancing accumulation of the latter^65–68^. However, the effect of envelope disruption on rifampicin activity appears to be more robust than its effect on azide-rifamycin accumulation (^13^, **Figure 3A**, and **Figure 4C-D, 4F**). A full account of this phenomenon is beyond the scope of the current work. However, our observation that the localization of cell wall metabolism changes in the presence of the drug, along with extensive literature reports of connections between rifampicin treatment and/or resistance and cell envelope synthesis and/or stress^8, 44–50^ suggest that synergy between inhibitors of transcription and envelope biosynthesis may be more complicated than permeation alone.

In contrast to bulk, cell-destroying methods for determining accumulation, *e.g*., mass spectrometry, we analyze accumulation in live cells. Our ability to track different cell populations by flow cytometry enabled a second point of nuance, namely that mycomembrane restriction is not only differential, both for fluorophores and antibiotics, but also heterogenous, for a subset of the same. Heterogeneity in *M. smegmatis* growth characteristics has been linked to differential susceptibility to cell envelope-acting drugs and to rifampicin^15, 16^. Further, more uniform cell growth rate in the absence of divisome factor LamA has been associated with enhanced killing by vancomycin and rifampicin in *M. smegmatis* and *M. tuberculosis*^14^. LamA was originally identified in a screen for mutants with more uniform uptake of a fluorescent indicator dye. These data suggest that mycomembrane permeation is one source of heterogeneity that contributes to antibiotic sensitivity.

We find that moxifloxacin accumulates in the periplasm less than ciprofloxacin, in agreement with its greater restriction by the mycomembrane (**Figures 4C-D** and **6B-C**). Given that moxifloxacin and ciprofloxacin inhibit DNA gyrase to similar extents *in vitro*^56^, other investigators have hypothesized that the enhanced permeation of moxifloxacin across the mycomembrane might explain its superior whole cell activity relative to other, less-hydrophobic fluoroquinolones^53^. Our data instead suggest that moxifloxacin may be more effective than ciprofloxacin despite being restricted to a greater extent by the mycomembrane. We speculate that moxifloxacin is more resistant to efflux or to xenobiotic metabolism than ciprofloxacin and/or is better than ciprofloxacin at traversing the inner membrane. The fluoroquinolone data suggest that the relationship between the hydrophobicity of a molecule and its accumulation (and activity, in the case of antibiotics) is not straightforward.

It is yet unclear how periplasm accumulation relates to whole cell accumulation measurements made by others, or to accumulation specifically in the cytoplasmic compartment. Novobiocin, rifampicin, and/or vancomycin have poor whole cell accumulation in *E. coli*, *Pseudomonas aeruginosa, M. abscessus*, and *M. tuberculosis* relative to other antibiotics tested^39–41, 69, 70^ and we find that azide derivatives of the cytoplasm-acting drugs likewise accumulate poorly in the mycobacterial periplasm. However, in contrast to the high periplasm accumulation that we observe for linezolid and tedizolid, these oxazolidinones accumulate poorly in whole *M. abscessus* and *P. aeruginosa* cells, likely because they are subject to efflux across the inner membrane^41,70^. Because we covalently trap azide-modified molecules in the cell wall, and because there is no known direct mechanism for efflux from the periplasm across the mycomembrane^71^, we speculate that PAC-MAN is less subject to efflux than whole cell measurements. Indeed, a strength of our current approach is that a periplasm-specific readout for accumulation enables us to assess the effect of mycomembrane perturbation close to the source. However, in the future, pairing cytoplasm-specific and periplasm-specific readouts for accumulation with defined perturbations to efflux, the inner membrane, and the mycomembrane will enable us to discern the relative contributions of different factors to molecule accumulation.

In this work we assayed *M. smegmatis* and *M. tuberculosis* under standard culture conditions to permit facile comparisons to the literature. However, we note that one of the culture medium ingredients, Tween-80 detergent, is known to strip away non-covalently bound lipids from the mycomembrane, notably phthiocerol dimycocerosate (PDIM)^72^. PDIM contributes to the envelope permeability barrier and restricts vancomycin activity in *M. tuberculosis*^72–74^. Beyond PDIM, it is well-known that environmental changes can result in concomitant changes to envelope structure, permeability, and/or antibiotic activity^5, 50, 62, 65, 69, 75^. By varying culture conditions, in the future we hope to untangle the contribution of *in vivo*-relevant environmental regulation to molecule accumulation in mycobacteria.

While here we have focused on the permeation of existing antibiotics, the molecule- and cell-specific sieving properties of the mycomembrane are important considerations in developing new, more effective drug regimens for tuberculosis. Strategies to promote antibiotic accumulation in Gram-negative species have generally fallen into two categories: disruption of the outer membrane, to non-specifically enhance drug permeation, or chemical modification of individual drugs, to promote permeation on a case-by-case basis. We suggest that the former strategy would not actually be non-specific in *M. tuberculosis*. Attention to the accumulation properties of individual drug candidates in individual cells may be necessary to optimize whole cell activity on a population-wide basis.

## Acknowledgments

We thank Dr. Joel Freundlich for valuable discussion, the director of the University of Massachusetts Amherst Flow Cytometry facilities, Dr. Amy Burnside, and the director of the University of Massachusetts Amherst Microscopy facilities, Dr. James Chambers for the help and advice. These studies were supported by the National Institutes of Health (R01 AI179080 to M.M.P. and M.S.S.) and T32 GM139789 (Chemistry Biology Interface Program at the University of Massachusetts Amherst, support for K.J.). L.P. was supported by ISPRO-Istituto per lo Studio, la Prevenzione e la Rete Oncologica, AIRC-Associazione Italiana Ricerca sul Cancro (IG #25694) and European Union (Next Generation EU, Mission 4, Component 2, CUP B93D21010860004, Spoke 6-RNA drug development).

## Materials and Methods

### Bacterial strains and growth conditions

*M. tuberculosis* mc^2^6206 (H37Rv Δ*panCD* Δ*leuCD*^76^) was provided by Dr. William Jacobs and was cultured in Middlebrook 7H9 (liquid) or on 7H10 (agar) supplemented with 0.5% (vol/vol) glycerol, 10% Middlebrook Oleic Albumin Dextrose Catalase (OADC; BD), and either 0.05% (vol/vol) Tyloxapol (Sigma), for strain maintenance, or 0.05% (vol/vol) Tween-80 (Sigma), for experimental manipulation. Growth media were additionally supplemented with 50 μg/ml L-leucine (Sigma) and 50 µg/ml pantothenic acid (Sigma). Wild-type, Δ*fbpA*, and complemented Δ*fbpA M. smegmatis* mc^2^155^77, 78^ were provided by Dr. Anil Ojha and cultured in Middlebrook 7H9 supplemented with 0.5% (vol/vol) glycerol, 1x ADC (10x ADC: 5 g bovine serum albumin, 2 g dextrose, 3 mg catalase in 100 mL), and 0.05% (vol/vol) Tween-80 (Sigma). Wild-type and *lptD*4213 E. coli K-12 MG1655 ^36–38^ were provided by Dr. Natacha Ruiz and grown in Lysogeny Broth (LB). All bacteria were grown 37° C shaking.

### Azide fluorophore labeling in diderm bacteria

*E. coli, M. smegmatis,* and *M. tuberculosis* were incubated with BCNDala^21^ probe (WuXi AppTec) at 37°C for 6 generations (*M. smegmatis* and *E. coli*) or 1 generation (*M. tuberculosis*). Bacteria were then washed twice with phosphate saline buffer (PBS) supplemented with 0.05% Tween-80 (PBST). *E. coli* and *M. smegmatis* cells were divided into aliquots then immediately reacted with 20 *μ*M of various azide-fluorophores (**Supplemental Figure 1** and **Supplemental Table 1**) for 1 hour at 37°C. *M. tuberculosis* was first treated with ethambutol (EMB) 100 *μ*g/mL overnight, washed to remove the EMB, then reacted with 20 *μ*M of azide-fluorophores. After azide-fluorophore reaction, bacteria were washed twice with PBST containing 0.01% BSA then fixed (*E. coli* and *M. smegmatis*: 2% formaldehyde for 15 min; *M. tuberculosis*: 4% paraformaldehyde for 30-120 minutes). Bacteria were washed again with PBST and analyzed by BD DUAL LSRFortessa flow cytometer or with Attune^TM^ NxT Acoustic Focusing Cytometer.

### PAC-MAN in mycobacteria (general)

Bacteria were grown to mid-log phase, diluted, and incubated with 25 *μ*M **TetD** probe^17^(WuXi AppTec). *M. smegmatis* was incubated in probe for 1-2 days to a final optical density (OD_600_) of 0.4. *M. tuberculosis* was incubated in probe for 4 days to a final OD_600_ of 0.2-0.3. Bacteria were washed twice with PBST (*M. smegmatis*) or 7H9 (*M. tuberculosis*) then incubated with 50 *μ*M of various test azides (usually azide antibiotics) in PBST (*M. smegmatis*) or 7H9 (*M. tuberculosis*) in 96-well plates for 2 hours at 37 °C shaking. Test azides were removed by spinning and supernatant removal and bacteria were then incubated with 10 *μ*M (*M. smegmatis*) or 17 *μ*M (*M. tuberculosis*) of the fluorogenic label AzCalfluor647 (**Supplemental Figure 1** and **Supplemental Table 1**) for 1 hour at 37 °C shaking. Bacteria were washed, fixed as above, then analyzed by BD DUAL LSRFortessa. Competition fluorescence was calculated as before^17^. First, fluorescence values of no-**TetD** controls (**Figure 4A**, step 1 omitted) were subtracted from all other samples. The adjusted fluorescence values were then divided by fluorescence values from no-test azide control bacteria (**Figure 4A**, step 2 omitted).

For **Supplemental Figure 3A-D**, to each well of a 96-well plate was added 50 *μ*L of DBCO-modified *M. smegmatis* followed by 50 *μ*L PBS containing different amounts of test azide to make up the indicated concentrations. Each test azide assayed in technical triplicate. The plates were incubated in 37 ℃ for 2 h. Bacteria were spun down for 2 min at 2700 g. The supernatant was decanted, and 100 *μ*L 50 *μ*M Az-6-FAM^17^ was then added to each well, followed by incubation in 37 ℃ for 1 h. Bacteria were then spun down again, and the pellets were washed twice with 100 *μ*L PBST then fixed with 100 *μ*L 4% formaldehyde for 15 min. Samples were analyzed by Attune^TM^ NxT Acoustic Focusing Cytometer.

### PAC-MAN in ethambutol-treated M. tuberculosis

*M. tuberculosis* was incubated with 25 *μ*M **TetD** for 72 hours to a final OD_600_ of 0.15, washed twice in 7H9, and treated +/− 100 *μ*g/mL ethambutol for 24 h. Ethambutol was removed by spinning and supernatant removal and bacteria were then incubated with the test azides as above.

### Antibiotic killing

Bacteria cultures were diluted to a final OD_600_ of 0.05. *M. tuberculosis* was treated with 100 *μ*g/mL EMB for 24 hours then spun at 4000 rpm and washed. Following this, 0.2 mL aliquots were added to each well of a 96 well plate. Antibiotics were added to each well to a concentration of 50 *μ*M to match the concentration of azide antibiotics used in PAC-MAN. The plates were incubated at 37 °C for 24 h. 20 *μ*L of 10-fold serial dilutions (in 7H9) were then plated on 7H10 agar. Agar plates were incubated at 37 °C and counted after 3 days (*M. smegmatis*) or 12-14 days (*M. tuberculosis*)

### DBCO-modified polystyrene beads and competition curves

DBCO-modified polystyrene beads were prepared as before^17, 42^. Briefly, 1 mL amino functionalized polystyrene beads (5% w/v, 5 mg) were spun down at 21,000 g for 15 min and washed with 1 mL deionized water before use. The beads were then spun down and resuspended in 10 mL 20 mM sodium borate buffer pH 9 with 1 *μ*g/mL DBCO-NHS ester and reacted in 37°C for 2 h with shaking. The resulting beads were spun down at 4000 g for 10 min, washed once and resuspended in sodium borate buffer. 200 *μ*L acetic anhydride was added to the suspension and reacted in 37°C for 2h with shaking. The resulting product was then spun down and washed twice with 10 mL PBS and resuspend in 20 mL PBS for storage.

DBCO-modified polystyrene beads were diluted 1:20 in PBS before adding to the assay. To each well of a MultiScreen HTS 96-well filter plate with 0.65 mm hydrophilic PVDF membrane were added 100 *μ*L beads. Beads were then filtered under vacuum to get the pellet of beads on the filter membrane. 50 *μ*M test azides, including azide antibiotics, were added to the bead’s pellets in each well with a final volume of 100 *μ*L, which were resuspended with volumetric pipettes. The plates were incubated in 37 ℃ for 2 h and then washed with 200 *μ*L PBS, filtered under vacuum to remove the supernatant. 100 *μ*L 50 *μ*M Az-6–FAM was then added to each well. The plates were then incubated in 37°C for 1h. The beads were washed with 200 *μ*L PBS twice by vacuum filtration and then resuspended in 200 *μ*L PBS. The samples were then analyzed by Attune^TM^ NxT Acoustic Focusing Cytometer.

In some cases, DBCO-modified polystyrene beads were subjected to the same protocol described above for *M. smegmatis* and analyzed by BD DUAL LSRFortessa. Results of test azide competitions were concordant between protocols.

### BODIPY-vancomycin labeling

*M. smegmatis* and *M. tuberculosis* were grown to OD_600_ 0.2 then incubated +/− EMB as above for one generation (3 hours for *M. smegmatis* and 24 hours for *M. tuberculosis*). EMB was removed and bacteria were then incubated with different concentrations of BODIPY-vancomycin (Thermo Fisher; mixed 1:1 with native vancomycin) for 6 hours in PBST for *M. smegmatis* or 7H9 for *M. tuberculosis*. Bacteria were then fixed as above and analyzed by flow cytometry.

### Cell wall labeling of rifampicin-treated M. smegmatis

*M. smegmatis* was grown until mid-log-phase then incubated +/− 10 *μ*g/mL rifampicin. Pulse-chase peptidoglycan labeling was performed by exposing bacteria to either 25 *μ*M 5-carboxytetramethylrhodamine-D-alanine (TADA/RADA^18, 27^, WuXi AppTec) or 1 mM alkyne-D-alanine (EDA/alkDA^18, 22^, Acros Organics) for 5 minutes at 37°C. Bacteria were centrifuged 5000 x g for 5 minutes and washed twice with pre-warmed 7H9. Following the washes, the bacteria were incubated for 15 minutes with pre-warmed 7H9 +/− 10 *μ*g/mL rifampicin at 37°C. After this incubation, 1 mM alkDA or 25 *μ*M RADA was added for 5 minutes. Samples were transferred to a chilled centrifuge and spun 5000 x g for 5 minutes then washed three times with cold PBSTB. Bacteria were fixed with 2% formaldehyde for 10 minutes at room temperature, followed by centrifugation at 5000 x g for 5 minutes and two washes with PBSTB. After fixing, the bacteria were subjected to copper-catalyzed alkyne-azide cycloaddition (CuAAC) reaction (200 *μ*M CuSO_4_, 800 *μ*M TBTA, 2.5 mM sodium ascorbate, 25 *μ*M AF488PicolylAz) for 60 min at room temperature. After CuAAC, the samples were centrifuged 5000 x g for 5 minutes and washed three times with PBSTB.

### Fluorescence microscopy

Fixed bacteria were imaged either by conventional fluorescence microscopy (Zeiss Axioscope A1 with 100x objectives), followed by analysis as described in ^27^, or by structured illumination microscopy (Nikon SIM-E/A1R with SR Apo TIRF 100x objective).

## Supplemental material summary

**Supporting Information:** supplemental figures (PDF).

## References

(1) Organization, W. H. Global Tuberculosis Report; 2021.

(2) Jarlier, V.; Nikaido, H. Mycobacterial cell wall: structure and role in natural resistance to antibiotics. FEMS Microbiol Lett 1994, 123 (1-2), 11–18. DOI: 10.1111/j.1574-6968.1994.tb07194.x From NLM Medline.

(3) Nikaido, H.; Jarlier, V. Permeability of the mycobacterial cell wall. Res Microbiol 1991, 142 (4), 437–443. DOI: 10.1016/0923-2508(91)90117-s From NLM Medline.

(4) Brennan, P. J.; Nikaido, H. The envelope of mycobacteria. Annu Rev Biochem 1995, 64, 29–63. DOI: 10.1146/annurev.bi.64.070195.000333 From NLM Medline.

(5) Dulberger, C. L.; Rubin, E. J.; Boutte, C. C. The mycobacterial cell envelope - a moving target. Nat Rev Microbiol 2020, 18 (1), 47–59. DOI: 10.1038/s41579-019-0273-7 From NLM Medline.

(6) Fullam, E.; Young, R. J. Physicochemical properties and. RSC Med Chem 2020, 12 (1), 43–56. DOI: 10.1039/d0md00265h.

(7) Vilcheze, C.; Molle, V.; Carrere-Kremer, S.; Leiba, J.; Mourey, L.; Shenai, S.; Baronian, G.; Tufariello, J.; Hartman, T.; Veyron-Churlet, R.;, et al. Phosphorylation of KasB regulates virulence and acid-fastness in Mycobacterium tuberculosis. PLoS Pathog 2014, 10 (5), e1004115. DOI: 10.1371/journal.ppat.1004115 From NLM Medline.

(8) Xu, W.; DeJesus, M. A.; Rucker, N.; Engelhart, C. A.; Wright, M. G.; Healy, C.; Lin, K.; Wang, R.; Park, S. W.; Ioerger, T. R.;, et al. Chemical Genetic Interaction Profiling Reveals Determinants of Intrinsic Antibiotic Resistance in Mycobacterium tuberculosis. Antimicrob Agents Chemother 2017, 61 (12). DOI: 10.1128/AAC.01334-17 From NLM Medline.

(9) Gao, L. Y.; Laval, F.; Lawson, E. H.; Groger, R. K.; Woodruff, A.; Morisaki, J. H.; Cox, J. S.; Daffe, M.; Brown, E. J. Requirement for kasB in Mycobacterium mycolic acid biosynthesis, cell wall impermeability and intracellular survival: implications for therapy. Mol Microbiol 2003, 49 (6), 1547–1563. DOI: 10.1046/j.1365-2958.2003.03667.x From NLM Medline.

(10) Philalay, J. S.; Palermo, C. O.; Hauge, K. A.; Rustad, T. R.; Cangelosi, G. A. Genes required for intrinsic multidrug resistance in Mycobacterium avium. Antimicrob Agents Chemother 2004, 48 (9), 3412–3418. DOI: 10.1128/AAC.48.9.3412-3418.2004 From NLM Medline.

(11) Liu, J.; Nikaido, H. A mutant of Mycobacterium smegmatis defective in the biosynthesis of mycolic acids accumulates meromycolates. Proc Natl Acad Sci U S A 1999, 96 (7), 4011–4016. DOI: 10.1073/pnas.96.7.4011 From NLM Medline.

(12) Nguyen, L.; Chinnapapagari, S.; Thompson, C. J. FbpA-Dependent biosynthesis of trehalose dimycolate is required for the intrinsic multidrug resistance, cell wall structure, and colonial morphology of Mycobacterium smegmatis. J Bacteriol 2005, 187 (19), 6603–6611. DOI: 10.1128/JB.187.19.6603-6611.2005 From NLM Medline.

(13) Li, S.; Poulton, N. C.; Chang, J. S.; Azadian, Z. A.; DeJesus, M. A.; Ruecker, N.; Zimmerman, M. D.; Eckartt, K. A.; Bosch, B.; Engelhart, C. A.;, et al. CRISPRi chemical genetics and comparative genomics identify genes mediating drug potency in Mycobacterium tuberculosis. Nat Microbiol 2022, 7 (6), 766–779. DOI: 10.1038/s41564-022-01130-y From NLM Medline.

(14) Rego, E. H.; Audette, R. E.; Rubin, E. J. Deletion of a mycobacterial divisome factor collapses single-cell phenotypic heterogeneity. Nature 2017, 546 (7656), 153–157. DOI: 10.1038/nature22361 From NLM Medline.

(15) Aldridge, B. B.; Fernandez-Suarez, M.; Heller, D.; Ambravaneswaran, V.; Irimia, D.; Toner, M.; Fortune, S. M. Asymmetry and aging of mycobacterial cells lead to variable growth and antibiotic susceptibility. Science 2012, 335 (6064), 100–104. DOI: 10.1126/science.1216166 From NLM Medline.

(16) Smith, T. C., 2nd; Pullen, K. M.; Olson, M. C.; McNellis, M. E.; Richardson, I.; Hu, S.; Larkins-Ford, J.; Wang, X.; Freundlich, J. S.; Ando, D. M.; et al. Morphological profiling of tubercle bacilli identifies drug pathways of action. Proc Natl Acad Sci U S A 2020, 117 (31), 18744–18753. DOI: 10.1073/pnas.2002738117 From NLM Medline.

(17) Liu, Z.; Lepori, I.; Chordia, M. D.; Dalesandro, B. E.; Guo, T.; Dong, J.; Siegrist, M. S.; Pires, M. M. A Metabolic-Tag-Based Method for Assessing the Permeation of Small Molecules Across the Mycomembrane in Live Mycobacteria. Angew Chem Int Ed Engl 2023, 62 (20), e202217777. DOI: 10.1002/anie.202217777 From NLM Medline.

(18) Kuru, E.; Hughes, H. V.; Brown, P. J.; Hall, E.; Tekkam, S.; Cava, F.; de Pedro, M. A.; Brun, Y. V.; VanNieuwenhze, M. S. In Situ probing of newly synthesized peptidoglycan in live bacteria with fluorescent D-amino acids. Angew Chem Int Ed Engl 2012, 51 (50), 12519–12523. DOI: 10.1002/anie.201206749 From NLM Medline.

(19) Liechti, G. W.; Kuru, E.; Hall, E.; Kalinda, A.; Brun, Y. V.; VanNieuwenhze, M.; Maurelli, A. T. A new metabolic cell-wall labelling method reveals peptidoglycan in Chlamydia trachomatis. Nature 2014, 506 (7489), 507–510. DOI: 10.1038/nature12892 From NLM Medline.

(20) Williams, M. A.; Aliashkevich, A.; Krol, E.; Kuru, E.; Bouchier, J. M.; Rittichier, J.; Brun, Y. V.; VanNieuwenhze, M. S.; Becker, A.; Cava, F.;, et al. Unipolar Peptidoglycan Synthesis in the Rhizobiales Requires an Essential Class A Penicillin-Binding Protein. mBio 2021, 12 (5), e0234621. DOI: 10.1128/mBio.02346-21 From NLM Medline.

(21) Shieh, P.; Siegrist, M. S.; Cullen, A. J.; Bertozzi, C. R. Imaging bacterial peptidoglycan with near-infrared fluorogenic azide probes. Proc Natl Acad Sci U S A 2014, 111 (15), 5456–5461. DOI: 10.1073/pnas.1322727111 From NLM Medline.

(22) Siegrist, M. S.; Whiteside, S.; Jewett, J. C.; Aditham, A.; Cava, F.; Bertozzi, C. R. (D)-Amino acid chemical reporters reveal peptidoglycan dynamics of an intracellular pathogen. ACS Chem Biol 2013, 8 (3), 500–505. DOI: 10.1021/cb3004995 From NLM Medline.

(23) Brown, A. R.; Gordon, R. A.; Hyland, S. N.; Siegrist, M. S.; Grimes, C. L. Chemical Biology Tools for Examining the Bacterial Cell Wall. Cell Chem Biol 2020, 27 (8), 1052–1062. DOI: 10.1016/j.chembiol.2020.07.024 From NLM Medline.

(24) Pidgeon, S. E.; Apostolos, A. J.; Nelson, J. M.; Shaku, M.; Rimal, B.; Islam, M. N.; Crick, D. C.; Kim, S. J.; Pavelka, M. S.; Kana, B. D.;, et al. L,D-Transpeptidase Specific Probe Reveals Spatial Activity of Peptidoglycan Cross-Linking. ACS Chem Biol 2019, 14 (10), 2185–2196. DOI: 10.1021/acschembio.9b00427 From NLM Medline.

(25) Lebar, M. D.; May, J. M.; Meeske, A. J.; Leiman, S. A.; Lupoli, T. J.; Tsukamoto, H.; Losick, R.; Rudner, D. Z.; Walker, S.; Kahne, D. Reconstitution of peptidoglycan cross-linking leads to improved fluorescent probes of cell wall synthesis. J Am Chem Soc 2014, 136 (31), 10874–10877. DOI: 10.1021/ja505668f From NLM Medline.

(26) Hsu, Y. P.; Rittichier, J.; Kuru, E.; Yablonowski, J.; Pasciak, E.; Tekkam, S.; Hall, E.; Murphy, B.; Lee, T. K.; Garner, E. C.;, et al. Full color palette of fluorescent d-amino acids for in situ labeling of bacterial cell walls. Chem Sci 2017, 8 (9), 6313–6321. DOI: 10.1039/c7sc01800b From NLM PubMed-not-MEDLINE.

(27) Garcia-Heredia, A.; Pohane, A. A.; Melzer, E. S.; Carr, C. R.; Fiolek, T. J.; Rundell, S. R.; Lim, H. C.; Wagner, J. C.; Morita, Y. S.; Swarts, B. M.;, et al. Peptidoglycan precursor synthesis along the sidewall of pole-growing mycobacteria. Elife 2018, 7. DOI: 10.7554/eLife.37243 From NLM Medline.

(28) Agard, N. J.; Prescher, J. A.; Bertozzi, C. R. A strain-promoted [3 + 2] azide-alkyne cycloaddition for covalent modification of biomolecules in living systems. J Am Chem Soc 2004, 126 (46), 15046–15047. DOI: 10.1021/ja044996f From NLM Medline.

(29) Sletten, E. M.; Bertozzi, C. R. Bioorthogonal chemistry: fishing for selectivity in a sea of functionality. Angew Chem Int Ed Engl 2009, 48 (38), 6974–6998. DOI: 10.1002/anie.200900942 From NLM Medline.

(30) Lepori I., O. Y., Im J., Ghosh N., Paul M., Schubert U.S., Fedeli S. Bioorthogonal “Click” Cycloadditions: A Toolkit for Modulating Polymers and Nanostructures in Living Systems. reactions 2024, 5 (1), 231–245.

(31) Goude, R.; Amin, A. G.; Chatterjee, D.; Parish, T. The arabinosyltransferase EmbC is inhibited by ethambutol in Mycobacterium tuberculosis. Antimicrob Agents Chemother 2009, 53 (10), 4138–4146. DOI: 10.1128/AAC.00162-09 From NLM Medline.

(32) McNeil, M. B.; Chettiar, S.; Awasthi, D.; Parish, T. Cell wall inhibitors increase the accumulation of rifampicin in Mycobacterium tuberculosis. Access Microbiol 2019, 1 (1), e000006. DOI: 10.1099/acmi.0.000006 From NLM PubMed-not-MEDLINE.

(33) Zhang, L.; Zhao, Y.; Gao, Y.; Wu, L.; Gao, R.; Zhang, Q.; Wang, Y.; Wu, C.; Wu, F.; Gurcha, S. S.;, et al. Structures of cell wall arabinosyltransferases with the anti-tuberculosis drug ethambutol. Science 2020, 368 (6496), 1211–1219. DOI: 10.1126/science.aba9102 From NLM Medline.

(34) Mikusova, K.; Slayden, R. A.; Besra, G. S.; Brennan, P. J. Biogenesis of the mycobacterial cell wall and the site of action of ethambutol. Antimicrob Agents Chemother 1995, 39 (11), 2484–2489. DOI: 10.1128/AAC.39.11.2484 From NLM Medline.

(35) Kilburn, J. O.; Takayama, K. Effects of ethambutol on accumulation and secretion of trehalose mycolates and free mycolic acid in Mycobacterium smegmatis. Antimicrob Agents Chemother 1981, 20 (3), 401–404. DOI: 10.1128/AAC.20.3.401 From NLM Medline.

(36) Sampson, B. A.; Misra, R.; Benson, S. A. Identification and characterization of a new gene of Escherichia coli K-12 involved in outer membrane permeability. Genetics 1989, 122 (3), 491–501. DOI: 10.1093/genetics/122.3.491 From NLM Medline.

(37) Braun, M.; Silhavy, T. J. Imp/OstA is required for cell envelope biogenesis in Escherichia coli. Mol Microbiol 2002, 45 (5), 1289–1302. DOI: 10.1046/j.1365-2958.2002.03091.x From NLM Medline.

(38) Ruiz, N.; Falcone, B.; Kahne, D.; Silhavy, T. J. Chemical conditionality: a genetic strategy to probe organelle assembly. Cell 2005, 121 (2), 307–317. DOI: 10.1016/j.cell.2005.02.014 From NLM Medline.

(39) Krishnamoorthy, G.; Leus, I. V.; Weeks, J. W.; Wolloscheck, D.; Rybenkov, V. V.; Zgurskaya, H. I. Synergy between Active Efflux and Outer Membrane Diffusion Defines Rules of Antibiotic Permeation into Gram-Negative Bacteria. mBio 2017, 8 (5). DOI: 10.1128/mBio.01172-17 From NLM Medline.

(40) Richter, M. F.; Drown, B. S.; Riley, A. P.; Garcia, A.; Shirai, T.; Svec, R. L.; Hergenrother, P. J. Predictive compound accumulation rules yield a broad-spectrum antibiotic. Nature 2017, 545 (7654), 299–304. DOI: 10.1038/nature22308 From NLM Medline.

(41) Geddes, E. J.; Gugger, M. K.; Garcia, A.; Chavez, M. G.; Lee, M. R.; Perlmutter, S. J.; Bieniossek, C.; Guasch, L.; Hergenrother, P. J. Porin-independent accumulation in Pseudomonas enables antibiotic discovery. Nature 2023, 624 (7990), 145–153. DOI: 10.1038/s41586-023-06760-8 From NLM Medline.

(42) Kelly, J. J.; Dalesandro, B. E.; Liu, Z.; Chordia, M. D.; Ongwae, G. M.; Pires, M. M. Measurement of Accumulation of Antibiotics to Staphylococcus aureus in Phagosomes of Live Macrophages. Angew Chem Int Ed Engl 2024, 63 (3), e202313870. DOI: 10.1002/anie.202313870 From NLM Medline.

(43) Piddock, L. J.; Williams, K. J.; Ricci, V. Accumulation of rifampicin by Mycobacterium aurum, Mycobacterium smegmatis and Mycobacterium tuberculosis. J Antimicrob Chemother 2000, 45 (2), 159–165. DOI: 10.1093/jac/45.2.159 From NLM Medline.

(44) Campodonico, V. L.; Rifat, D.; Chuang, Y. M.; Ioerger, T. R.; Karakousis, P. C. Altered Mycobacterium tuberculosis Cell Wall Metabolism and Physiology Associated With RpoB Mutation H526D. Front Microbiol 2018, 9, 494. DOI: 10.3389/fmicb.2018.00494 From NLM PubMed-not-MEDLINE.

(45) Giddey, A. D.; Ganief, T. A.; Ganief, N.; Koch, A.; Warner, D. F.; Soares, N. C.; Blackburn, J. M. Cell Wall Proteomics Reveal Phenotypic Adaption of Drug-Resistant Mycobacterium smegmatis to Subinhibitory Rifampicin Exposure. Front Med (Lausanne) 2021, 8, 723667. DOI: 10.3389/fmed.2021.723667 From NLM PubMed-not-MEDLINE.

46. Patel, Y.; Soni, V.; Rhee, K. Y.; Helmann, J. D. Mutations in rpoB That Confer Rifampicin Resistance Can Alter Levels of Peptidoglycan Precursors and Affect beta-Lactam Susceptibility. mBio 2023, 14 (2), e0316822. DOI: 10.1128/mbio.03168-22 From NLM Medline.

(47) Vinella, D.; D’Ari, R.; Jaffe, A.; Bouloc, P. Penicillin binding protein 2 is dispensable in Escherichia coli when ppGpp synthesis is induced. EMBO J 1992, 11 (4), 1493–1501. DOI: 10.1002/j.1460-2075.1992.tb05194.x From NLM Medline.

(48) du Preez, I.; Loots du, T. Altered fatty acid metabolism due to rifampicin-resistance conferring mutations in the rpoB Gene of Mycobacterium tuberculosis: mapping the potential of pharmaco-metabolomics for global health and personalized medicine. OMICS 2012, 16 (11), 596–603. DOI: 10.1089/omi.2012.0028 From NLM Medline.

(49) Sebastian, J.; Nair, R. R.; Swaminath, S.; Ajitkumar, P. Mycobacterium tuberculosis Cells Surviving in the Continued Presence of Bactericidal Concentrations of Rifampicin in vitro Develop Negatively Charged Thickened Capsular Outer Layer That Restricts Permeability to the Antibiotic. Front Microbiol 2020, 11, 554795. DOI: 10.3389/fmicb.2020.554795 From NLM PubMed-not-MEDLINE.

(50) Koh, E. I.; Oluoch, P. O.; Ruecker, N.; Proulx, M. K.; Soni, V.; Murphy, K. C.; Papavinasasundaram, K.; Reames, C. J.; Trujillo, C.; Zaveri, A.;, et al. Chemical-genetic interaction mapping links carbon metabolism and cell wall structure to tuberculosis drug efficacy. Proc Natl Acad Sci U S A 2022, 119 (15), e2201632119. DOI: 10.1073/pnas.2201632119 From NLM Medline.

(51) Haemers, A.; Leysen, D. C.; Bollaert, W.; Zhang, M. Q.; Pattyn, S. R. Influence of N substitution on antimycobacterial activity of ciprofloxacin. Antimicrob Agents Chemother 1990, 34 (3), 496–497. DOI: 10.1128/AAC.34.3.496 From NLM Medline.

(52) Govendir, M.; Hansen, T.; Kimble, B.; Norris, J. M.; Baral, R. M.; Wigney, D. I.; Gottlieb, S.; Malik, R. Susceptibility of rapidly growing mycobacteria isolated from cats and dogs, to ciprofloxacin, enrofloxacin and moxifloxacin. Vet Microbiol 2011, 147 (1-2), 113–118. DOI: 10.1016/j.vetmic.2010.06.011 From NLM Medline.

(53) Danilchanka, O.; Pavlenok, M.; Niederweis, M. Role of porins for uptake of antibiotics by Mycobacterium smegmatis. Antimicrob Agents Chemother 2008, 52 (9), 3127–3134. DOI: 10.1128/AAC.00239-08 From NLM Medline.

(54) Gillespie, S. H.; Billington, O. Activity of moxifloxacin against mycobacteria. J Antimicrob Chemother 1999, 44 (3), 393–395. DOI: 10.1093/jac/44.3.393 From NLM Medline.

(55) Sulochana, S.; Rahman, F.; Paramasivan, C. N. In vitro activity of fluoroquinolones against Mycobacterium tuberculosis. J Chemother 2005, 17 (2), 169–173. DOI: 10.1179/joc.2005.17.2.169 From NLM Medline.

(56) Aldred, K. J.; Blower, T. R.; Kerns, R. J.; Berger, J. M.; Osheroff, N. Fluoroquinolone interactions with Mycobacterium tuberculosis gyrase: Enhancing drug activity against wild-type and resistant gyrase. Proc Natl Acad Sci U S A 2016, 113 (7), E839–846. DOI: 10.1073/pnas.1525055113 From NLM Medline.

(57) Peraro, L.; Deprey, K. L.; Moser, M. K.; Zou, Z.; Ball, H. L.; Levine, B.; Kritzer, J. A. Cell Penetration Profiling Using the Chloroalkane Penetration Assay. J Am Chem Soc 2018, 140 (36), 11360–11369. DOI: 10.1021/jacs.8b06144 From NLM Medline.

(58) Sheldrick, G. M.; Jones, P. G.; Kennard, O.; Williams, D. H.; Smith, G. A. Structure of vancomycin and its complex with acetyl-D-alanyl-D-alanine. Nature 1978, 271 (5642), 223–225. DOI: 10.1038/271223a0 From NLM Medline.

(59) Nieto, M.; Perkins, H. R. Physicochemical properties of vancomycin and iodovancomycin and their complexes with diacetyl-L-lysyl-D-alanyl-D-alanine. Biochem J 1971, 123 (5), 773–787. DOI: 10.1042/bj1230773 From NLM Medline.

(60) Perkins, H. R. Specificity of combination between mucopeptide precursors and vancomycin or ristocetin. Biochem J 1969, 111 (2), 195–205. DOI: 10.1042/bj1110195 From NLM Medline.

(61) Cho, H.; Uehara, T.; Bernhardt, T. G. Beta-lactam antibiotics induce a lethal malfunctioning of the bacterial cell wall synthesis machinery. Cell 2014, 159 (6), 1300–1311. DOI: 10.1016/j.cell.2014.11.017 From NLM Medline.

(62) Pohane, A. A.; Carr, C. R.; Garhyan, J.; Swarts, B. M.; Siegrist, M. S. Trehalose Recycling Promotes Energy-Efficient Biosynthesis of the Mycobacterial Cell Envelope. mBio 2021, 12 (1). DOI: 10.1128/mBio.02801-20 From NLM Medline.

(63) Kohanski, M. A.; Dwyer, D. J.; Hayete, B.; Lawrence, C. A.; Collins, J. J. A common mechanism of cellular death induced by bactericidal antibiotics. Cell 2007, 130 (5), 797–810. DOI: 10.1016/j.cell.2007.06.049 From NLM Medline.

(64) Jewett, J. C.; Bertozzi, C. R. Cu-free click cycloaddition reactions in chemical biology. Chem Soc Rev 2010, 39 (4), 1272–1279. DOI: 10.1039/b901970g From NLM Medline.

(65) Poulton, N. C.; Rock, J. M. Unraveling the mechanisms of intrinsic drug resistance in Mycobacterium tuberculosis. Front Cell Infect Microbiol 2022, 12, 997283. DOI: 10.3389/fcimb.2022.997283 From NLM Medline.

(66) Hui, J.; Gordon, N.; Kajioka, R. Permeability barrier to rifampin in mycobacteria. Antimicrob Agents Chemother 1977, 11 (5), 773–779. DOI: 10.1128/AAC.11.5.773 From NLM Medline.

(67) Bellerose, M. M.; Proulx, M. K.; Smith, C. M.; Baker, R. E.; Ioerger, T. R.; Sassetti, C. M. Distinct Bacterial Pathways Influence the Efficacy of Antibiotics against Mycobacterium tuberculosis. mSystems 2020, 5 (4). DOI: 10.1128/mSystems.00396-20 From NLM PubMed-not-MEDLINE.

(68) Jakkala, K.; Ajitkumar, P. Hypoxic Non-replicating Persistent Mycobacterium tuberculosis Develops Thickened Outer Layer That Helps in Restricting Rifampicin Entry. Front Microbiol 2019, 10, 2339. DOI: 10.3389/fmicb.2019.02339 From NLM PubMed-not-MEDLINE.

(69) Sarathy, J.; Dartois, V.; Dick, T.; Gengenbacher, M. Reduced drug uptake in phenotypically resistant nutrient-starved nonreplicating Mycobacterium tuberculosis. Antimicrob Agents Chemother 2013, 57 (4), 1648–1653. DOI: 10.1128/AAC.02202-12 From NLM Medline.

(70) McGowen K., F. T., Wang X., Zinga S., Wolf I.D., Akusobi C.C., Denkinger C.M., Rubin E.J., Sullivan M.R. Efflux pumps and membrane permeability contribute to intrinsic antibiotic resistance in *Mycobacterium abscessus*. bioRxiv 2024, preprint. DOI: 10.1101/2024.08.23.609441.

(71) Remm, S.; Earp, J. C.; Dick, T.; Dartois, V.; Seeger, M. A. Critical discussion on drug efflux in Mycobacterium tuberculosis. FEMS Microbiol Rev 2022, 46 (1). DOI: 10.1093/femsre/fuab050 From NLM Medline.

(72) Mulholland, C. V.; Wiggins, T. J.; Cui, J.; Vilcheze, C.; Rajagopalan, S.; Shultis, M. W.; Reyes-Fernandez, E. Z.; Jacobs, W. R., Jr.; Berney, M. Propionate prevents loss of the PDIM virulence lipid in Mycobacterium tuberculosis. Nat Microbiol 2024, 9 (6), 1607–1618. DOI: 10.1038/s41564-024-01697-8 From NLM Medline.

(73) Camacho, L. R.; Constant, P.; Raynaud, C.; Laneelle, M. A.; Triccas, J. A.; Gicquel, B.; Daffe, M.; Guilhot, C. Analysis of the phthiocerol dimycocerosate locus of Mycobacterium tuberculosis. Evidence that this lipid is involved in the cell wall permeability barrier. J Biol Chem 2001, 276 (23), 19845–19854. DOI: 10.1074/jbc.M100662200 From NLM Medline.

(74) Wang, Q.; Boshoff, H. I. M.; Harrison, J. R.; Ray, P. C.; Green, S. R.; Wyatt, P. G.; Barry, C. E., 3rd. PE/PPE proteins mediate nutrient transport across the outer membrane of Mycobacterium tuberculosis. Science 2020, 367 (6482), 1147–1151. DOI: 10.1126/science.aav5912 From NLM Medline.

(75) Larrouy-Maumus, G.; Marino, L. B.; Madduri, A. V.; Ragan, T. J.; Hunt, D. M.; Bassano, L.; Gutierrez, M. G.; Moody, D. B.; Pavan, F. R.; de Carvalho, L. P. Cell-Envelope Remodeling as a Determinant of Phenotypic Antibacterial Tolerance in Mycobacterium tuberculosis. ACS Infect Dis 2016, 2 (5), 352–360. DOI: 10.1021/acsinfecdis.5b00148 From NLM PubMed-not-MEDLINE.

(76) Sampson, S. L.; Dascher, C. C.; Sambandamurthy, V. K.; Russell, R. G.; Jacobs, W. R., Jr.; Bloom, B. R.; Hondalus, M. K. Protection elicited by a double leucine and pantothenate auxotroph of Mycobacterium tuberculosis in guinea pigs. Infect Immun 2004, 72 (5), 3031–3037. DOI: 10.1128/IAI.72.5.3031-3037.2004 From NLM Medline.

(77) Snapper, S. B.; Melton, R. E.; Mustafa, S.; Kieser, T.; Jacobs, W. R., Jr. Isolation and characterization of efficient plasmid transformation mutants of Mycobacterium smegmatis. Mol Microbiol 1990, 4 (11), 1911–1919. DOI: 10.1111/j.1365-2958.1990.tb02040.x From NLM Medline.

(78) Ojha, A. K.; Trivelli, X.; Guerardel, Y.; Kremer, L.; Hatfull, G. F. Enzymatic hydrolysis of trehalose dimycolate releases free mycolic acids during mycobacterial growth in biofilms. J Biol Chem 2010, 285 (23), 17380–17389. DOI: 10.1074/jbc.M110.112813 From NLM Medline.

