## Supplemental file 1 for "The mycomembrane differentially and heterogeneously restricts antibiotic permeation"

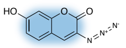

1. 3-Az-7-Coumarin

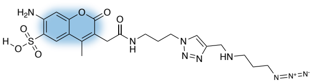

2. AFDye350-Azplus

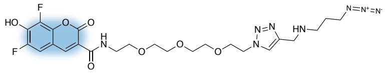

3. PB-Azplus

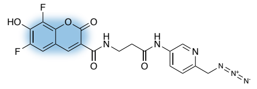

4. PB-picolylAz

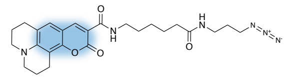

5. Az-Coumarin343X

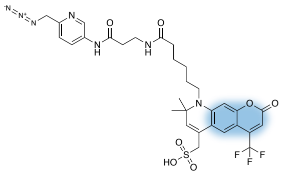

6. AFDye430-picolylAz

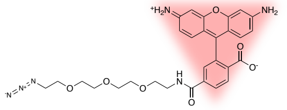

7. Az-Carboxyrhodamine110

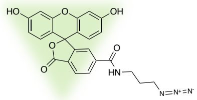

8. Az-6-FAM

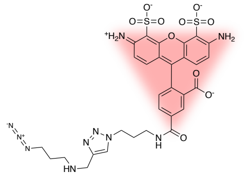

9. AFDye488Azplus

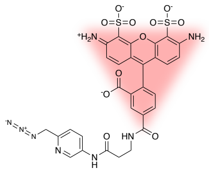

10. AF488PicolylAz

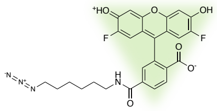

11. AzOG488

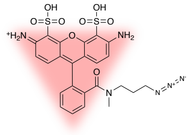

12. AzATTO488

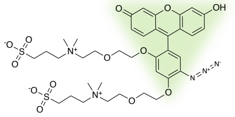

13. AzCalFluor488

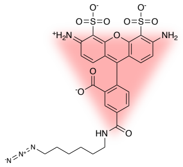

14. AzAFdye488

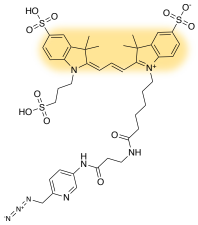

15. Cy3-PicolylAz

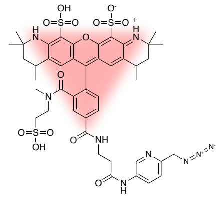

16. MB543PicolylAz

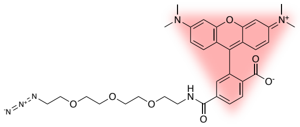

17. AzFluor545

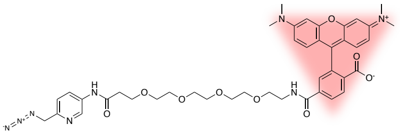

18. 5TAMRAPicolylAz

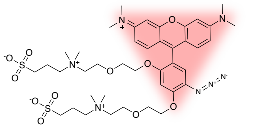

19. AzCalFluor555

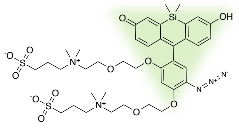

20. AzCalFluor580

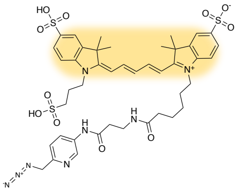

21. PicolylAzCy5

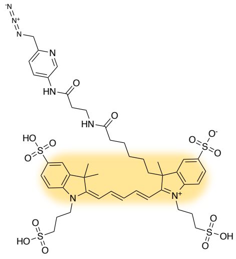

22. PicolylAz647AF

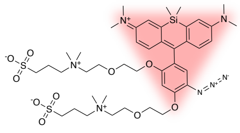

23. AzCalFluor647

**Supplemental Figure 1. Fluorophore chemical structure.** Azide fluorophores used in **Figure 1**. Dye families based on main scaffold are highlighted: coumarin (blue), rhodamine (red), fluorescein (green), and cyanine (yellow).

| Az-fluorophore # | Az-fluorophore name | Flow Cytometry fluorescent channel* | Commercial source |
|---|---|---|---|
| 1 | 3-Az-7-hydroxycoumarin | Indo-1 (Violet) | Sigma-Aldrich |
| 2 | AFdye350-Azplus | Indo-1 (Violet) | Vector Laboratories |
| 3 | PB-Azplus | BV421 | Vector Laboratories |
| 4 | PB-picolylAz | BV421 | Vector Laboratories |
| 5 | Az-Coumarin343X | BV510 | Lumiprobe |
| 6 | AF430-picolylAz | BV605 | Vector Laboratories |
| 7 | Az-Carboxyrhodamine110 | FITC | Vector Laboratories |
| 8 | Az-6-FAM | FITC | Lumiprobe |
| 9 | AF488Azplus | FITC | Vector Laboratories |
| 10 | AF488PiAz | FITC | Vector Laboratories |
| 11 | AzOG488 | FITC | Vector Laboratories |
| 12 | AzAtto488 | FITC | Atto-Tech |
| 13 | AzCalFluor488 | FITC | Vector Laboratories |
| 14 | AzAFdye488 | FITC | Vector Laboratories |
| 15 | Cy3-PicolylAz | PE-Texas Red | Vector Laboratories |
| 16 | MB543PicolylAz | PE-Texas Red | Vector Laboratories |
| 17 | AzFluor545 | PE-Texas Red | Sigma-Aldrich |
| 18 | 5TAMRAPicolylAz | PE-Texas Red | Vector Laboratories |
| 19 | AzCalFluor555 | PE-Texas Red | Vector Laboratories |
| 20 | AzCalFluor580 | PE-Texas Red | Vector Laboratories |
| 21 | Cy5-PicolylAz | APC | Vector Laboratories |
| 22 | PicolylAz647AF | APC | Vector Laboratories |
| 23 | AzCalFluor647 | APC | Vector Laboratories |

*fluorescent channels refer to **BD DUAL LSRFortessa**, 5 excitation lasers (355nm, 405nm, 488nm, 561nm, and 640nm), 18 color analysis capabilities. Optical configuration details: https://www.umass.edu/ials/sites/default/files/facilities/2021-Fortessa-Oplical-Configuration.pdf

**Supplemental Table 1. Azide fluorophore additional details.** Azide fluorophore (**Supplemental Figure 1**) numbers, names, fluorescent channels used for the flow cytometry analysis, and commercial sources.

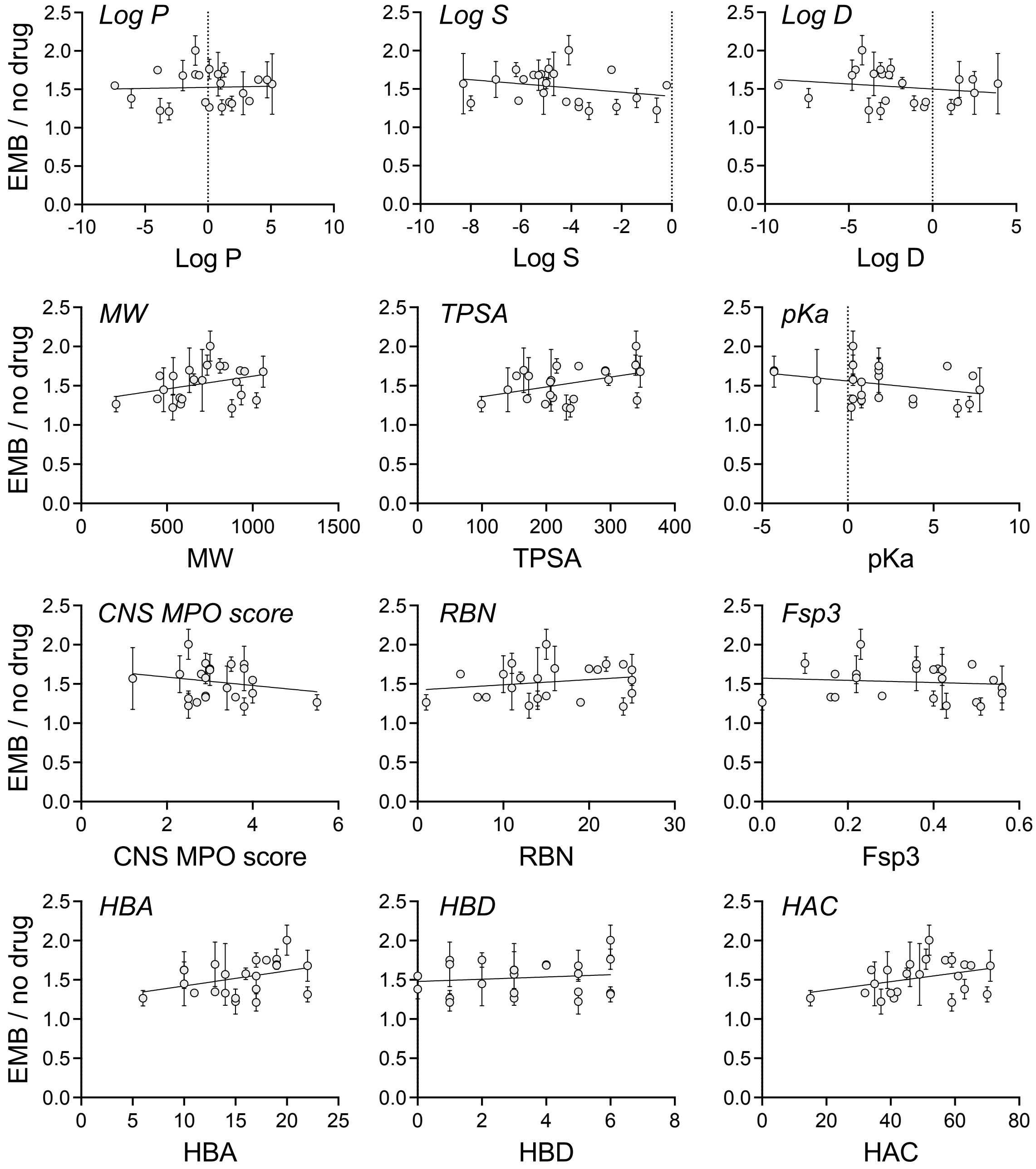


**Supplemental Figure 2. Lack of correlation between physicochemical properties of azide fluorophores and sensitivity to mycomembrane perturbation.** Physicochemical properties, calculated with Collaborative Drug Discovery (CDD) Vault software, of the azide fluorophores. The fluorescence ratios for *M. tuberculosis* ethambutol vs. no drug (**Figure 1E**) were plotted as function of each physicochemical property. Simple linear regression analyses did not identify obvious correlations between fluorophore sensitivity to mycomembrane perturbation and the investigated properties. Log P: *n*-Octanol/water partition coefficient (log); Log S: water solubility (log); Log D: *n*-Octanol/water distribution coefficient at pH 7.4 (log); MW: molecular weight; TPSA: topological polar surface area; pKa: acid dissociation constant (-log); CNS MPO score: central nervous system multiparameter optimization; RBN: rotatable bonds; Fps3: fraction of $sp^3$ carbons; HBA: hydrogen bond acceptor; HBD: hydrogen bond donor; HAC: heavy atom count.

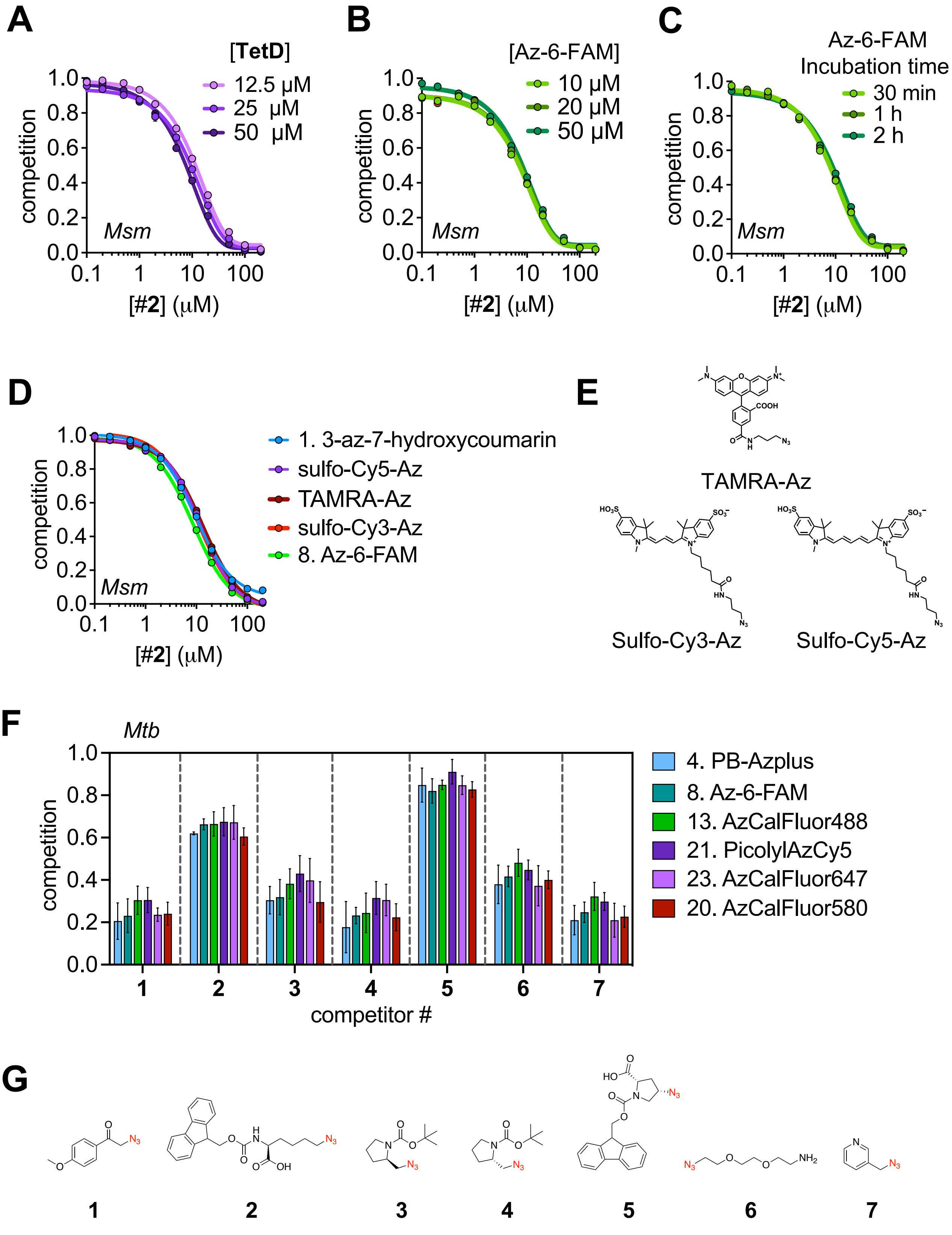


**Supplemental Figure 3. PAC-MAN assay performance over different experimental parameters.** See next page for full figure legend.

**Supplemental Figure 3. PAC-MAN assay performance over different experimental parameters.** PAC-MAN assay performance over different experimental parameters, including the concentration of **TetD** probe (A), the concentration of azide fluorophore (B), the incubation time of azide fluorophore (C), and the choice of azide fluorophore (D) and (F). (E) shows the structures of azide fluorophores not included in **Supplemental Figure 1** or **Supplemental Table 1** and (G) shows the structures of test azides used in (A)-(D) and (F). Competition levels of test azides in (F) are comparable between azide fluorophores but not to each other due to intrinsic differences in azide antibiotic reactivity. See **Figure 6** and accompanying text for details. Data in (F) are mean +/- SEM of three independent biological replicates and are not statistically significant. Data in (A)-(D) are mean +/- SEM for 3 technical replicates and representative of three independent biological replicates. *Msm: M. smegmatis; Mtb: M. tuberculosis.*

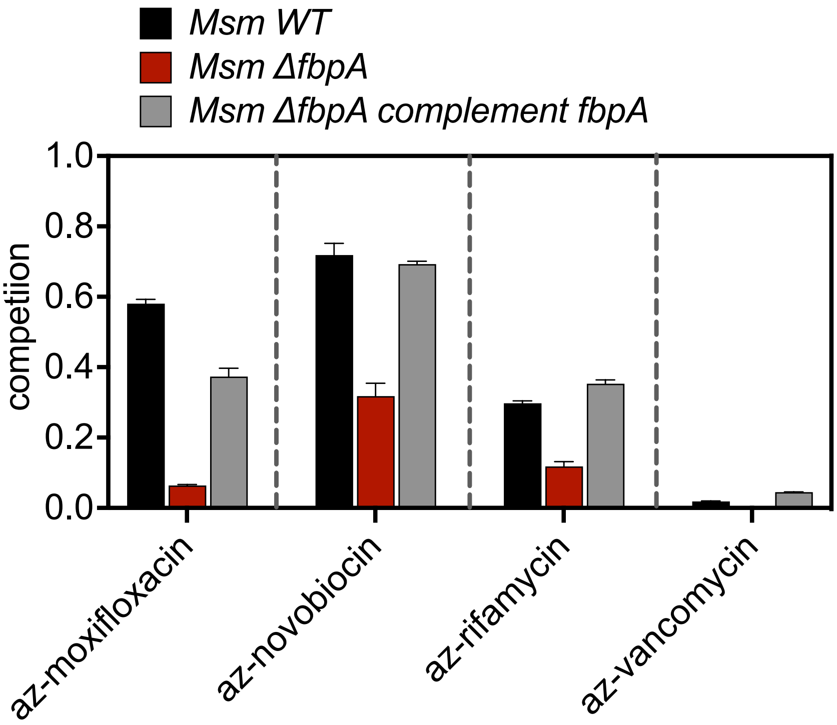


**Supplemental Figure 4.** ***M. smegmatis fbpA*** **complementation rescues competition phenotype.** PAC-MAN assay performed as in **Figure 4C** for *M. smegmatis* wild-type, Δ*fbpA*, and Δ*fbpA* complement. Competition levels of azide antibiotics are comparable between strains but not to each other due to intrinsic differences in azide- reactivity. See Figure 6 and accompanying text for details. Data are mean +/- SEM of 3 technical replicates.

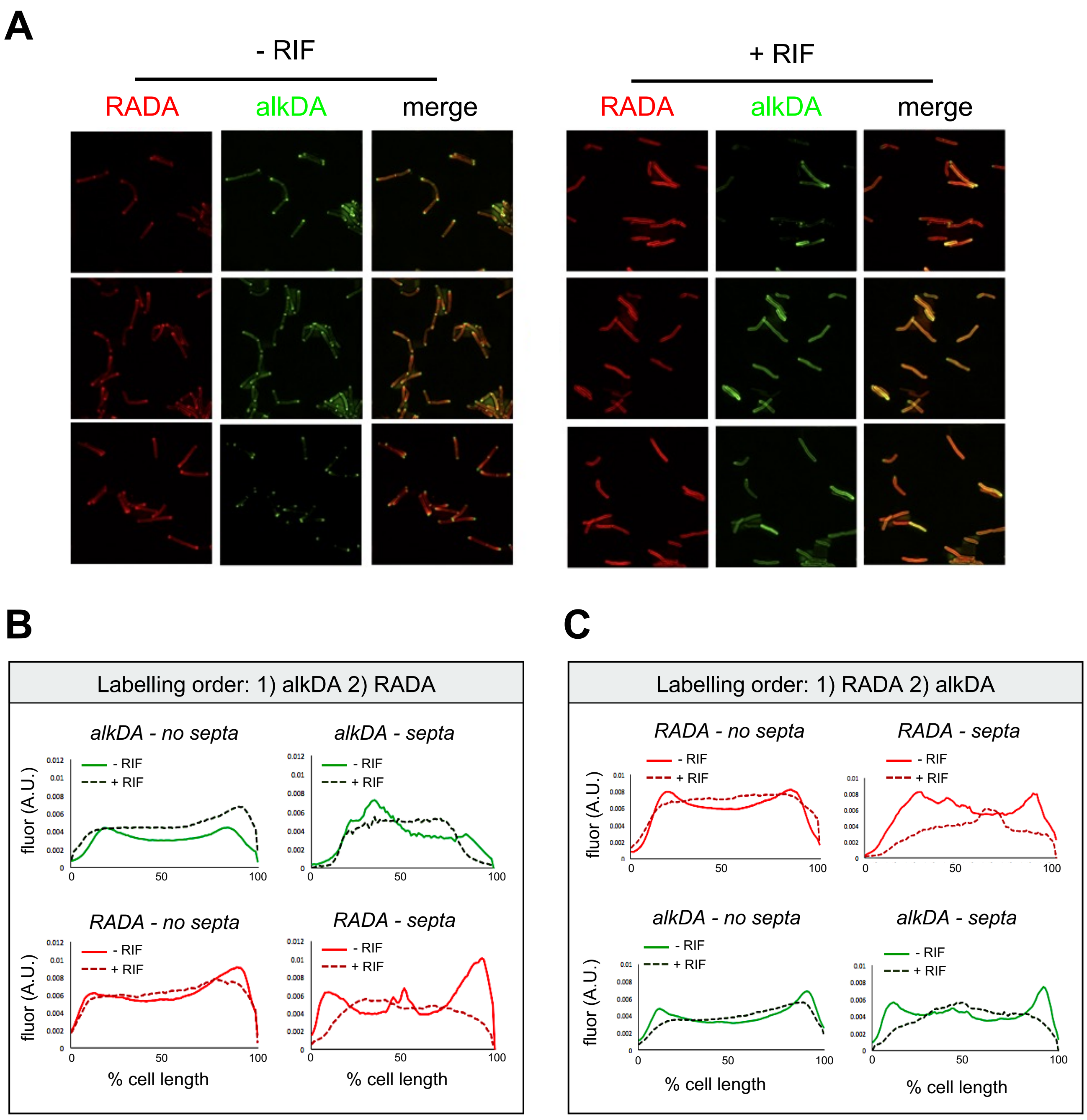


**Supplemental Figure 5. Rifampicin treatment delocalizes active cell wall metabolism.** See next page for full figure legend.

**Supplemental Figure 5. Rifampicin treatment delocalizes active cell wall metabolism.** *M. smegmatis* were treated +/- rifampicin then labeled with peptidoglycan cell wall probes 5-carboxytetramethylrhodamine-d-alanine (TADA/RADA[18, 27], red) or alkyne-d-alanine (EDA/alkDA[18, 22], green) for 5 minutes. Bacteria were washed, grown +/- rifampicin in the absence of probe for 15 minutes, then labeled again with the other probe for an additional 5 minutes. Bacteria were washed again then subjected to copper-catalyzed alkyne-azide cycloaddition (CuAAC) with AF488PicolylAz to reveal the presence of cell wall-embedded alkDA. (A) Structured illumination microscopy (SIME). (B-C) Quantification of the cell fluorescence (from images obtained via conventional light microscopy) for *M. smegmatis* incubated with alkDA followed by RADA (B) or with inverted order (C). Cellular fluorescence was quantitated[27] for n=81 (no drug; alkDA -> RADA), n=41 (rifampicin; alkDA -> RADA), n=140 (no drug; RADA -> alkDA), or n=58 (rifampicin; RADA -> alkDA). Fluorescence was normalized to cell length and total fluorescence intensity. Cells were oriented such that the brighter pole was on the right-hand side of the graph. In rifampicin-treated *M. smegmatis*, RADA and alkDA labeling is less pronounced than untreated at normal sites of active cell wall metabolism (elongating poles and dividing septa). A.U.: arbitrary units.

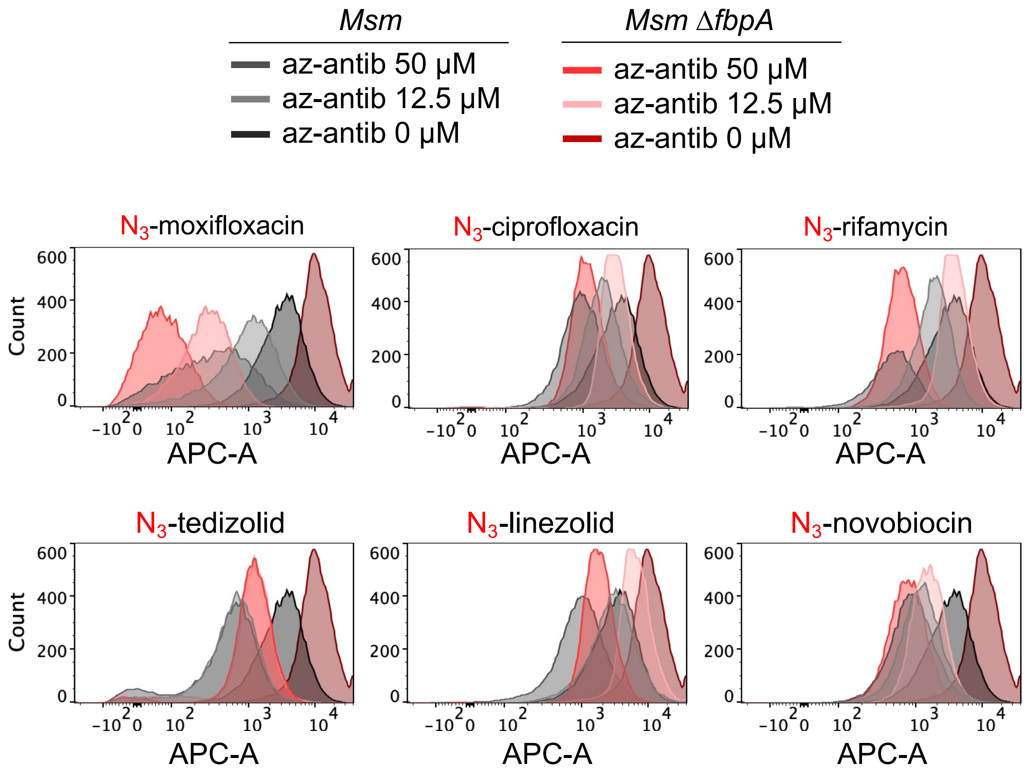


**Supplemental Figure 6. Restriction of non-vancomycin antibiotics by mycomembrane is more uniform than restriction of vancomycin.** Flow cytometry histograms from PAC-MAN for azide drugs other than vancomycin. In contrast to **Figure 5B**, we did not observe obvious, reproducible heterogeneity in competition from azide-moxifloxacin, -ciprofloxacin, -rifamycin, -tedizolid, -linezolid, or -novobiocin.
